# A quantitative biophysical principle to explain the 3D cellular connectivity in curved epithelia

**DOI:** 10.1101/2020.02.19.955567

**Authors:** Pedro Gómez-Gálvez, Pablo Vicente-Munuera, Samira Anbari, Antonio Tagua, Carmen Gordillo-Vázquez, Jesús A. Andrés-San Román, Daniel Franco-Barranco, Ana M. Palacios, Antonio Velasco, Carlos Capitán-Agudo, Clara Grima, Valentina Annese, Ignacio Arganda-Carreras, Rafael Robles, Alberto Márquez, Javier Buceta, Luis M. Escudero

## Abstract

Epithelial cell organization and the mechanical stability of tissues are closely related. In this context, it has been recently shown that packing optimization in bended/folded epithelia is achieved by a surface tension energy minimization mechanism that leads to a novel cellular shape: the *scutoid*. However, further cellular and tissue level implications of this new developmental paradigm remain unknown. Here we focus on the relationship between this complex cellular shape and the connectivity between cells. We address this problem using a combination of computational, experimental, and biophysical approaches in tubular epithelia. In particular, we examine how energy drivers affect the three-dimensional packing of these tissues. We challenge our biophysical model by reducing the cell adhesion in epithelial cells. As a result, we observed an increment on the cell apico-basal intercalation propensity that correlated with a decrease of the energy barrier necessary to connect with new cells. We conclude that tubular epithelia satisfy a quantitative biophysical principle, that links tissue geometry and energetics with the average cellular connectivity.

## INTRODUCTION

During the last decades much progress has been achieved in the understanding of the emergence of self-organization in tissues. This problem has been addressed from the viewpoint of energetics considerations (Alt et al., 2017; Canela-Xandri et al., 2011; Fletcher et al., 2014; Misra et al., 2017; Nelson et al., 2005; Siedlik et al., 2017; Sugimura et al., 2016; Trepat et al., 2009), material-like properties (Bi et al., 2015; Campàs et al., 2014; Latorre et al., 2018; Mongera et al., 2018; Pérez-González et al., 2019; Yang et al., 2017), and the analysis of the cellular packing (Curran et al., 2017; Farhadifar et al., 2007; Gibson et al., 2006; Gibson et al., 2011; Gómez et al., 2021; Honda, 1978; Lewis, 1928; Mao et al., 2013; Sanchez-Gutierrez et al., 2016; Thompson, 1945). As for the latter, the analysis of epithelial surfaces as tessellations of convex polygons has revealed mathematical and physical principles with biological consequences. One well-known example are the implications of the celebrated Euler’s formula, *V* – *E* + *F* = *χ* **(STAR Methods)** (Euler, 1767). This formula implies that cells in packed tissues have, on average, six neighbors (i.e., the average cellular connectivity on a surface reads 〈*n*_2*D*_〉 = 6) (Reinhardt, 1918; Wetzel, 1926). This principle has biological consequences, for example, the degree of cellular connectivity regulates the strength of the cell-cell juxtracrine signaling (Guignard et al., 2020; Sharma et al., 2019; Tung et al., 2012).

For a long time, the validity of this mathematical concept (i.e., each cell, on average, connects with six neighboring cells) has been assumed in three dimensions (3D): 〈*n*_2*D*_〉 = 6 = ⇒ 〈*n*_3*D*_〉 = 6. Such an assumption is rooted in the common idealization of epithelial cells as regular prismatic solids in either planar or bended epithelia. However, the recent discovery of more complex cellular shapes in epithelia, i.e., scutoids, that achieve an efficient 3D tissue packing has set a new paradigm that has not been yet fully explored **(Box A)** (Gómez-Gálvez et al., 2018; Mughal et al., 2018; Rupprecht et al., 2017). Scutoidal cellular shapes are the result of intercalations among cells along the apico-basal axis **(Box A-C** and **Fig. 1A).** This phenomenon is then a spatial version of the so-called T1 transitions that produce rearrangements of neighboring cells in the plane as a function of time in numerous developmental processes **(Box B)** (Bertet et al., 2004; Irvine and Wieschaus, 1994; Spencer et al., 2017). Importantly, scutoids imply necessarily changes in the neighboring relationship between cells in a 3D spatial context and, consequently, modify the connectivity properties of cells **(Box C).** Still, the analysis of tissue organization in 3D and the corresponding biophysical insight have been hindered by the technical difficulties to accurately segment and 3D-reconstruct cells, especially in curved tissues. In addition, very few computational models account for the presence of apico-basal transitions to investigate 3D self-organization in tissues (Gómez-Gálvez et al., 2018; Ioannou et al., 2020; Mughal et al., 2018; Okuda et al., 2019; Rupprecht et al., 2017). Moreover, from an energetics viewpoint, while the appearance of scutoids can be explained by a minimal model based on a surface/line tension minimization mechanism (Gómez-Gálvez et al., 2018; Mughal et al., 2018; Okuda et al., 2019; Rupprecht et al., 2017), the role played by additional energetic contributions to modulate the frequency of apico-basal intercalations is unknown.

**Figure 1.**
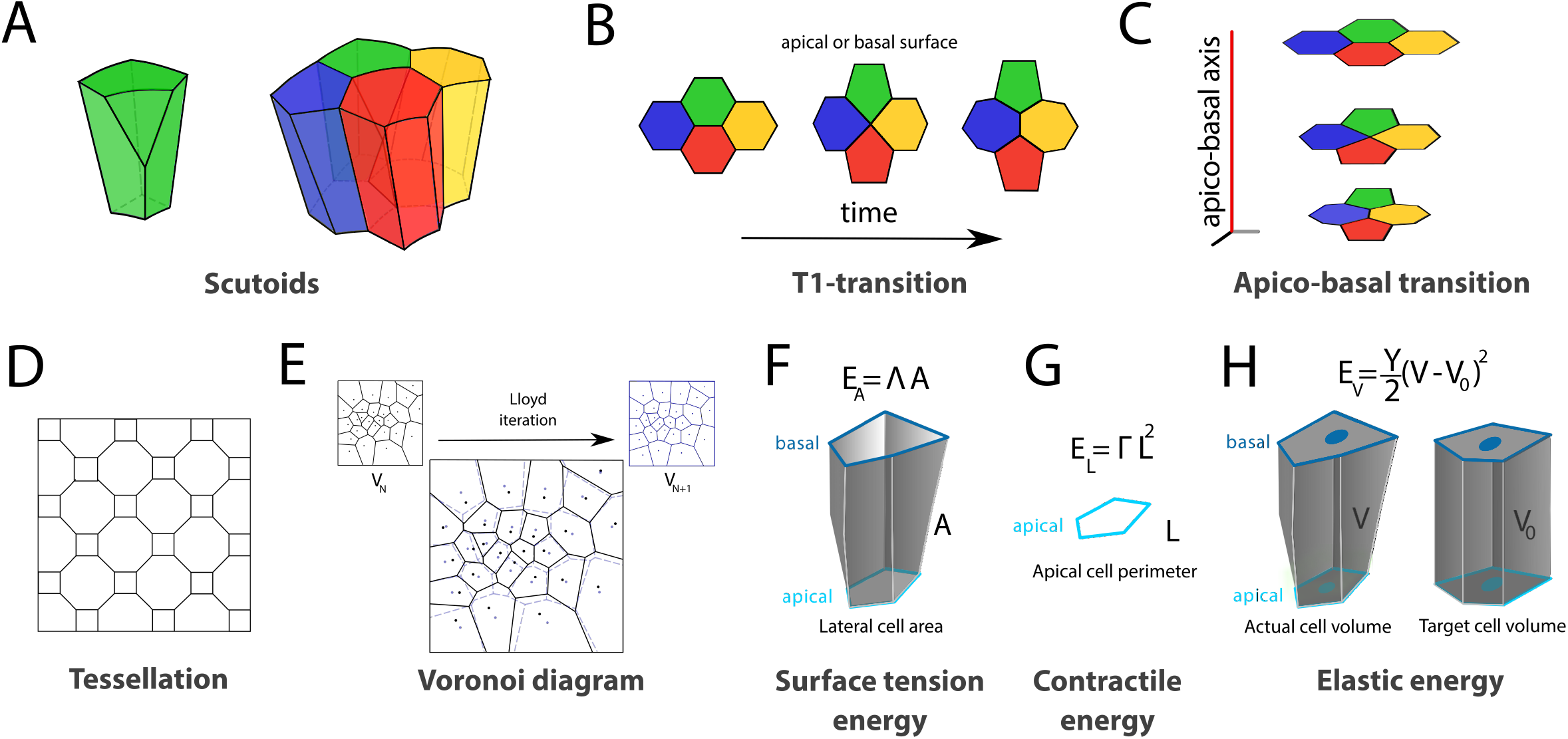
*In silico* Voronoi epithelial model: energetics and analysis of apico-basal cell intercalations in tubes. **A:** (left) Scutoids entail apico-basal intercalations among packing cells that can be envisioned as *spatial* T1 transitions to exchange neighbors (right): the green and the red cells are neighbors in the basal surface (but not in the apical surface) while the opposite is true for the blue and the yellow cells. **B:** *In silico* Voronoi 3D epithelial models are generated by populating with cell seeds (circles) the apical surface, ∑_*a*_. The location of cell seeds at any other surface/plane, *∑,* is obtained by implementing apical normal projections, ***n***, up to the basal surface, *∑_b_*. At each surface, *∑,* a 2D Voronoi tessellation is performed and the 3D cellular shape of the cell is built upon the collection of these tessellations. **C:** (top) In the particular case of tubular epithelia, normal projections of apical cell seeds correspond to radial projections, and the thickness/curvature of tubes are characterized by the surface ratio, *s* = *R/R_a_* (apical/basal surfaces: light/dark blue. *R*: dashed yellow). (bottom) Illustrative rendering of a Voronoi tube cell. **D:** The so-called CVT scale (iterations of Lloyd’s algorithm, **STAR Methods)** measures the topological disorder in *in silico* tubular epithelia and leads to different cellular morphologies. In a V1 (Voronoi 1) model cell seeds are randomly distributed on the apical surface to generate a planar Voronoi tessellation **(STAR Methods).** By applying the Lloyd algorithm iteratively, the apical topological disorder diminishes (top to bottom: V1, V5, and V10 examples). The random location of seeds in V1 implies the emergence of a wide range of different polygonal cell types. As Lloyd’s algorithm iterates, the larger the tessellation order is **(Box E).** We observe this progressive ordering from a V1 to V10 as polygonal distributions converge to results with a larger proportion of hexagons and a reduction of the other polygonal shapes (inset: polygon distribution insets for V1, V5, and V10). The average number of apico-basal intercalations per cell in *in silico* tubes (*n* = 20 realizations per CVT scale, each tube composed by 200 cells) increases with the tissue thickness (surface ratio) but does not change significantly with the CVT scale. The black/red/green arrows correspond to the illustrative examples of the planar apical tessellations shown on the left. **E:** The average lateral cell area as a function of the CVT and the surface ratio (values normalized to the V1 case at *s_b_* = 10, *n* = 20 per CVT scale) indicates that the average surface tension energy does not depend on the level of topological disorder. **F:** Average (green bar) and standard deviation (red lines) of the apical cellular perimeter as a function of the CVT scale. Values normalized with respect to the V1 case (*n* = 20 per CVT scale). **G**: The cellular volume variance is a proxy for the average elastic cell energy **(STAR Methods).** The latter decreases with the CVT scale and increases with the surface ratio (tissue thickness) (*n* = 20 per CVT scale). Volume values were normalized with respect to the V1 case with 〈*V*(*s_b_* = 10)〉 = 1. **H:** Cross-correlation between average energy profiles along the apico-basal axis, 〈*E_A_*〉 (dark grey), 〈*E_V_*〉 (red), and 〈*E_A_*〉 + 〈*E_L_*〉 (light grey), and the number of apico-basal intercalations, 〈*i*〉. Solid lines stand for the averages among disorder configurations (i.e., CVT scale) and the dotted lines delimit the standard deviation band. 〈*E_V_*〉 + 〈*E_L_*〉 has not been plotted since the extra contribution of the contractile term, 〈*E_L_*〉, does not modify the correlation function. **I:** Cross-correlation between energy gradients (*∂_S_*〈*E_A_*〉 and *∂_S_*〈*E_V_*〉) and the gradient of intercalations along the apico-basal axis (*∂_s_*〈*i*〉). Color code as in H.

The analysis of 3D packing is in turn utterly relevant in cubic and columnar monolayer tubular epithelia, where scutoids appear more frequently (Gómez-Gálvez et al., 2018; Gómez et al., 2021; Iruela-Arispe and Beitel, 2013; Sanchez-Corrales et al., 2018). Epithelial tubes are in fact the primary developmental structures in all organisms with bilateral symmetry (Gilbert and Barresi, 2013), and tubulogenesis is fundamental in a broad variety of key developmental processes, including gastrulation and neurulation (Colas and Schoenwolf, 2001; Iruela-Arispe and Beitel, 2013; Leptin and Grunewald, 1990; Nelson, 2009; Pilot and Lecuit, 2005; Röper, 2018; Swanson and Beitel, 2006). Furthermore, epithelial tubes are the essential functional unit of many mammalian organs, including glands, components of the digestive apparatus, lungs, and kidney (Huebner and Ewald, 2014).

Here, we study the packing and the 3D cellular connectivity properties of epithelial tubes. We analyze the effect of different energetic contributions to modulate the frequency of apico-basal intercalations; demonstrate that the presence of scutoids implies a breakdown of the principle 〈*n*_3*D*_〉 = 6; and reveal a quantitative biophysical principle that links the 3D cellular connectivity, energetics, and geometrical descriptors (e.g., tissue curvature/thickness). Our findings are supported by i) a computational model that realistically renders the 3D cellular organization of tubular epithelia (including the appearance of scutoids); ii) experimental data of wildtype (wt) and mutant epithelial tubes (*Drosophila*’s salivary gland) whose 3D cellular structure has been accurately characterized by means of a computer-aided image analysis pipeline. And iii), a biophysical model, supported by mathematical calculations, that connects the tissue energetics with the 3D organization of epithelial tubes.

## RESULTS

### The Voronoi computational tubular model supports a relationship between energy profiles and the intercalation propensity

To understand how the geometry of tubular epithelia and different energy contributions affect the 3D cellular packing and connectivity of these tissues, we designed and implemented a computational epithelial model that follows the principles of Voronoi tessellations **(Box D, E)** (Gómez-Gálvez et al., 2018). In brief, we generated 3D models of epithelial tubes by populating with seeds the apical surface, ∑_*α*_ (light blue points in **Fig. 1B)** and implementing normal projections of those seeds up to the basal surface, ∑_*b*_ (dark blue points in **Fig. 1B).** Each seed and its projection corresponded to an individual cell of a tube. At each surface section ∑ (from apical to basal) a 2D Voronoi diagram was performed, and the collection of those tessellations rendered the 3D cellular geometry of cells (see details in **STAR Methods).** We point out that we do not implement any temporal dynamics to the seeds. Thus, our computational model is suited to static epithelial configurations as the ones experimentally reported herein (see below).

Epithelial tubes appear in nature with very different thicknesses and cellular arrangements. In order to explore how these features influence the 3D packing properties of tubular epithelia we built diverse *in silico* Voronoi tubes. First, to investigate the effect of tissue thickness we computed Voronoi tubes with different surface ratios *s* = *R/R_a_* (*R* and *R_a_* being the radial coordinate of the tube and the apical radius respectively, **Fig. 1C, D).** We used *s*-steps of 0.5 up to *s* = 10, so we were able to explore 19 different values of the basal radius, *R_b_* **(Fig. 1D).** Second, we generated 10 different configurations in terms of the disorder level of the spatial positions of the cellular seeds on the apical surface and the corresponding Voronoi tessellations (V1 to V10, **Fig. 1D).** To that end, we used a fully random Voronoi tessellation, i.e., randomized positions of cellular seeds, as the most disordered pattern (V1). That configuration was made progressively more uniform (i.e., spatially ordered) after nine successive iterations of the homogenizing Lloyd’s algorithm **(Box E** and **STAR Methods) (Fig. 1D).** The resulting set of 10 different cellular arrangements (V1 to V10) with increasing order properties conforms a Centroidal Voronoi Tessellation (CVT) scale that has been proved useful to analyze the effect of the topological organization of tissues and to simulate different tissues and/or pathological conditions (Sanchez-Gutierrez et al., 2016; Vicente-Munuera et al., 2020). We used the CVT scale to investigate how the average number of apico-basal intercalations per cell, 〈*i*〉, changes as a function of the apico-basal coordinate, *s,* and the disorder level **(Fig. 1D).** As previously reported, we found that the number of apico-basal transitions **(Fig. 1D)** and scutoids **(Fig. S1)** increased with *s* (Gómez-Gálvez et al., 2018). As for the effect of the disorder level, we found that only in the case of fully disordered tubes (i.e., V1: random case), and for low values of *s*, there are more intercalations, whereas for the rest of cases we observed that 〈*i*〉 is fairly independent of the CVT scale **(Fig. 1D).**

Energy contributions can be linked to geometric features of the shapes of epithelial cells (Alt et al., 2017), see **Box F-H.** We used the set of Voronoi tubes (V1 to V10) to explore surface tension, elasticity, and apical contractility energies, since these energy contributions have been shown to play key roles in the organization of epithelia (Alt et al., 2017; Farhadifar et al., 2007). As a first step, we estimated the average cellular energy profiles as a function of s in the computational tubular model **(Fig. 1E-G).** The average surface tension energy **(Box F)** is related to the average lateral area of the cells, 〈*A*〉, and therefore increase with the surface ratio, *s.* Our results revealed that 〈*A*〉 is seemingly independent of the CVT scale **(Fig. 1E).** Consequently, the average cell surface tension energy profile does not depend on the level of the topological disorder. The contractile energy **(Box G)** is related to the average and the variance of the apical perimeter, *L,* (Gilbert and Barresi, 2013; Farhadifar et al., 2007) therefore it does not depend on the surface ratio, *s.* The Voronoi model revealed that 〈*L*〉 is CVT independent, but the apical perimeter fluctuations decrease as the CVT scale increases **(Fig. 1F).** Finally, the average cell elastic energy (Gelbart et al., 2012; Odell et al., 1981) depends on the average and the variance of the cellular volume **(Box H).** Since the average cell volume 〈*V*〉 is, by construction, independent of the CVT scale **(STAR Methods),** the average cellular elastic energy increases with the cellular volume fluctuations, that in turn decrease with the CVT scale **(Fig. 1G).**

In order to evaluate how the appearance of scutoids is modulated by these energy contributions for different values of the tissue thickness, we computed the cross-correlation functions, *C*(*s*), between the average cellular energy profiles, 〈*E_Z_*〉 (*Z* being *A* or *V,* i.e., surface tension or elastic terms), and the average number of apico-basal intercalations, 〈*i*〉 **(Fig. 1H** and **STAR Methods).** The cross-correlation measures the similarity between two signals as a function of the displacement (or lag) of one signal relative to the other. In our case the displacement/lag refers to the apico-basal coordinate, *s,* and consequently we inquire into the possibility that energetic contributions either precede or follow the appearance of apico-basal intercalations. Our results indicate that maximum correlations are obtained at zero lag independently of the disorder level and that the appearance of scutoids correlates more significantly with the surface tension energy profile than with the elastic energy: 95% vs. 80% respectively. The latter is in agreement with previous studies that have shown that surface tension energy minimization is the main cause underlying the appearance of scutoids (Gómez-Gálvez et al., 2018; Gómez et al., 2021; Mughal et al., 2018). Also, when assessing the extra effect of the energy input due to the apical contractility term to the surface tension energy, 〈*E_A_*〉 + 〈*E_L_*〉, we found that it increases the correlation between energy profile and the number of intercalations up to 98%, but it does not lead to any change in the correlation due to elastic terms, 〈*E_V_*〉 + 〈*E_L_*〉 **(Fig. 1H).**

We further examined the cross-correlation between the gradient of cellular intercalations along the apico-basal axis, *∂_s_*〈*i*〉 = *∂*〈*i*〉/*∂s*, and the gradient of the energy, *∂*_s_〈*E_Z_*〉. In this way, we evaluated the level of correspondence between the variation of the number of intercalations and the changes of the energy as a function of the radial coordinate, *s*. We found that, independently of the CVT scale, *∂_s_*〈*i*〉 correlates slightly stronger with changes in the surface tension energy, *∂_s_*〈*E_A_*〉, than with changes of the elastic contribution, *∂_s_*〈*E_V_*〉: ~80% versus ~75% respectively at optimal lag **(Fig. 1I).** Interestingly, *∂_s_*〈*i*〉 lags behind *∂_s_*〈*E_Z_*〉, i.e., the optimal lag for which *C*(*s*) is the largest is located at *s* > 0. Therefore, energy variations along the apico-basal axis seem to precede changes in the number of intercalations that, in turn, suggests an instructive role of the former over the latter.

Summing up, the Voronoi tubular model supports the idea that surface tension energy is the more relevant contribution regulating the appearance of apico-basal intercalations, and suggests that elastic terms play a role, yet less important than surface tension, for modulating the intercalation propensity (see **Discussion).**

### The Voronoi tubular model suggests a link between 3D tissue packing and energy cues

To link quantitatively energy traits and 3D packing, we implemented a benchmark able to simultaneously reveal the existence of apico-basal intercalations (scutoids) and the polygonal distribution of cells at the apical and the basal surfaces. To that end, we computed the probability that cells change their polygonal class between the apical and basal surfaces. Thus, the components (i.e., bins) of this distribution along the diagonal account for cells that have the same polygonal class at apical and basal surfaces, whereas the spreading away from the diagonal ensures the existence of scutoids and, consequently, changes in the cellular 3D connectivity **(Fig. 2A** and **STAR Methods).** Our data revealed that, regardless of the value of the tissue thickness (and the CVT scale), the dominant apical-basal polygonal class corresponds to cells with six neighbors **(Fig. 2A).** As the tissue thickness, *s_b_* = *R_b_/R_a_*, increases, more scutoidal shapes with a distinct number of neighbors in apical and basal surfaces appeared. This feature was revealed by the increasing value of the spreading away from the diagonal, *η*^2^ **(STAR Methods** and **Table S1).** In that regard, in agreement with the results shown in **Fig. 1D-G,** our data indicates that *η*^2^ increases with the tissue thickness and decreases with the CVT scale **(Fig. 2B).** Also, the cross-correlation analysis between the spreading coefficient and energy profiles agrees with **Fig. 1H** and reveals that independently of the CVT scale the neighbor exchanges correlate more strongly with the surface tension energy profile than with the elastic contribution: 90% vs. 70% at zero (optimal) lag **(Fig. 2C).** Furthermore, we computed the average number of total contacts between cells (i.e., the average 3D cellular connectivity), 〈*n*_3*D*_〉, as a function of the surface ratio (i.e., the radial coordinate) and the Voronoi class (i.e., the level of cellular disorder in the tissue) **(Fig. S1).** Our data indicated, that cells, on average, are connected to more than six cells, i.e., 〈*n*_3*D*_〉 > 6, and the results are quantitatively consistent with a mathematical derivation that shows that 〈*n*_3*D*_〉 is linearly proportional to the number of apico-basal intercalations 〈*i*〉: 〈*n*_3*D*_〉 = 6 + 〈*i*〉/2 **(STAR Methods** and **Fig. S1).**

**Figure 2.**
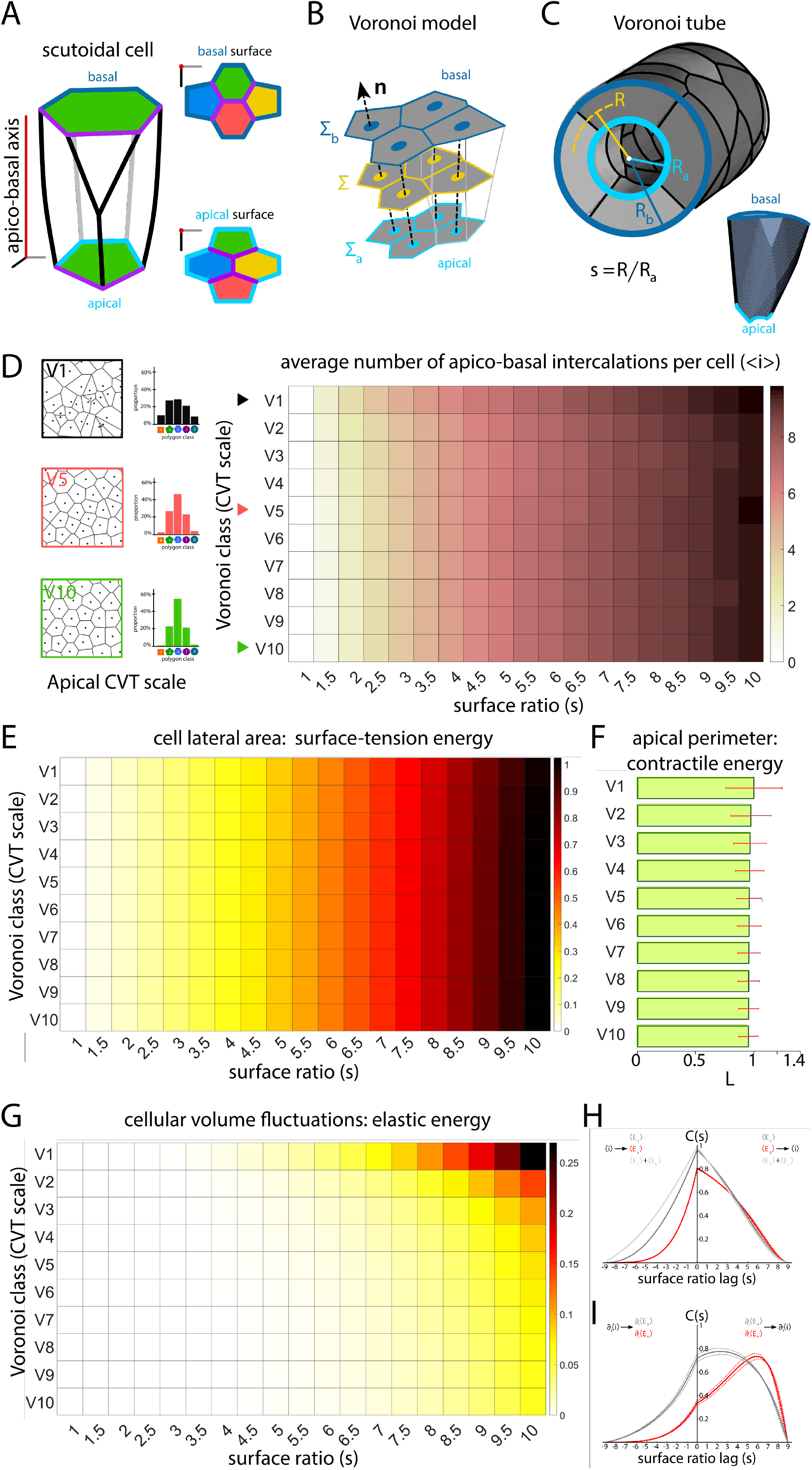
Three-dimensional packing and connectivity properties of the Voronoi tubular model. **A:** (left) Schematic representation of a 3D histogram that accounts for the probability that cells have *n_a_* (number) of neighbors in the apical surface and *n_b_* neighbors in the basal surface. Cells with the same polygonal class in apical and basal surfaces contribute to bins along the diagonal (red squares). The bins spreading away from the diagonal (green squares) ensures the presence of scutoidal cells. E.g.: the red and green cells shown in the plot contribute to the bins indicated in the 3D histogram (red and green stars respectively). (right) 3D histograms of V5 tubes for increasing values of the surface ratio. A larger value of the spreading coefficient, *η^2^,* **(STAR Methods)** indicates an increasing number of scutoids. **B:** Density plot showing the value of the spreading coefficient, *η^2^,* of 3D histograms as a function of the surface ratio and the Voronoi class in *in silico* tubes (*n* = 20 tubes per CVT scale). **C:** Cross-correlation between average energy profiles along the apico-basal axis, 〈*E_A_*〉 (dark grey), 〈*E_V_*〉 (red), and 〈*E_A_*〉 + 〈*E_L_*〉 (light grey), and the spreading coefficient, *η*^2^. Solid lines stand for the averages among disorder configurations (i.e., CVT scale) and the dotted lines delimit the standard deviation band. 〈*E_V_*〉 + 〈*E_L_*〉 has not been plotted since the extra contribution of the contractile term, 〈*E_L_*〉, does not modify the correlation function.

### The Voronoi tubular model recapitulates the properties of *in vivo* epithelial tubes

In order to compare the results obtained in our Voronoi computational tubular models against the properties found in real tissues, we implemented a methodological pipeline that combines several image analysis techniques to accurately reconstruct the 3D shapes of cells of *in vivo* epithelial tubes (Arganda-Carreras et al., 2017; Franco-Barranco et al., 2021; Machado et al., 2019) **(STAR Methods).** We used the *Drosophila* larval salivary gland, a cubic monolayer epithelium, as a model due to its ideal characteristics to study complex tubular developmental structures (Girdler and Roper, 2014) **(Fig. 3A).** Also, cellular growth and division, as well as possible global tissue deformation processes, do not occur in the *Drosophila*’s salivary gland at the developmental stage of our observations (i.e. the tissue is static); a fact that enables the comparison with the Voronoi computational model.

**Figure 3.**
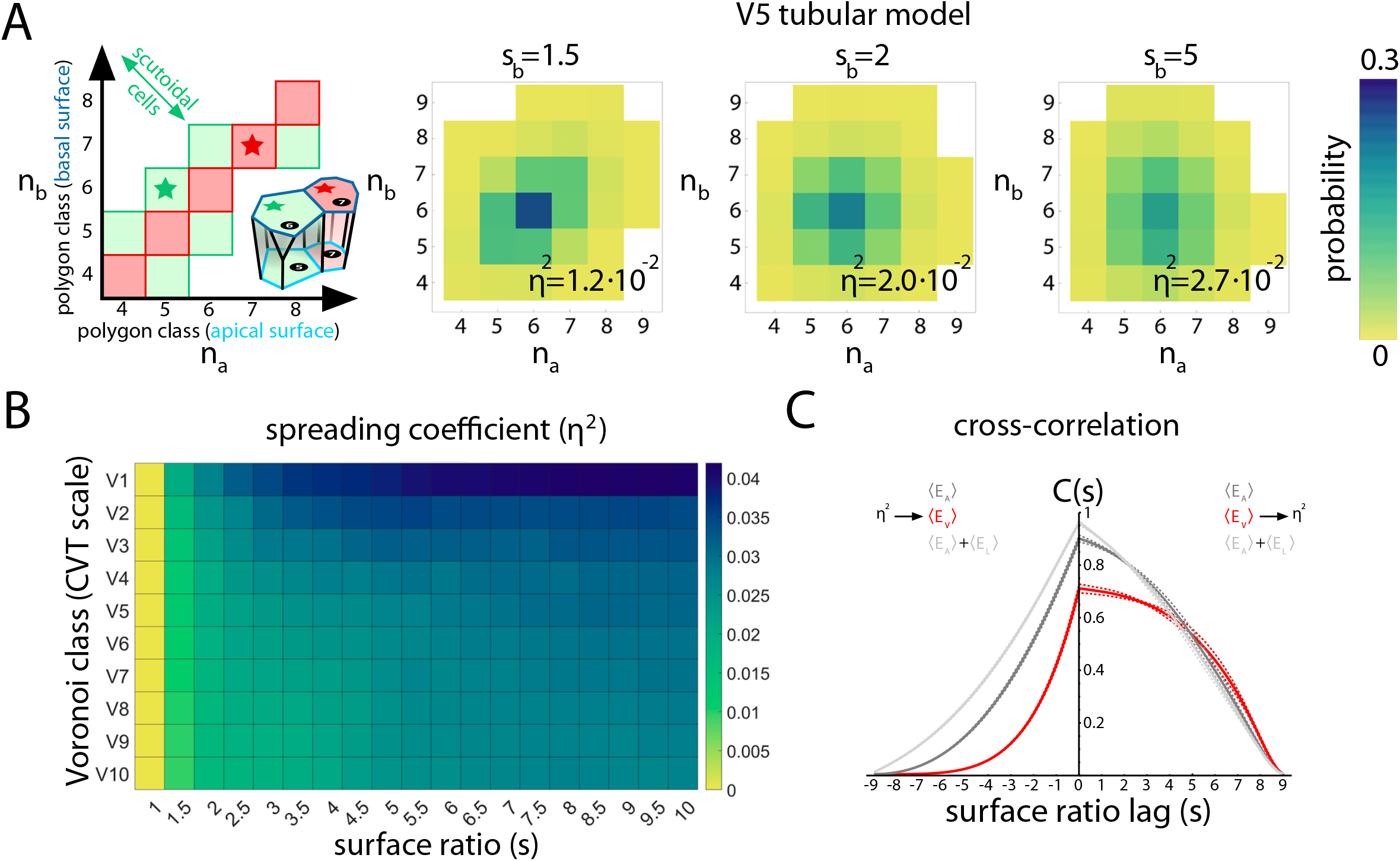
*Drosophila’s* salivary gland analysis. **A:** (top) Full projection of a wildtype salivary gland (cell contours stained by Cy3-labeled phalloidin, **STAR Methods).** (middle) Computer rendering of the segmented salivary gland shown on top. Scale bar 100*μm*. (bottom) 3D rendering of a representative segmented salivary gland. **B:** Density plot of the 3D distribution of neighbor exchanges between apical and basal surfaces as a function of the number of neighbors in apical, *n_a_,* and basal, *n_b_,* surfaces (as in **Fig. 2A)** in wildtype salivary glands (*n* = 20 glands, 979 cells). **C:** Average profiles of the number of apico-basal intercalations, 〈*i*〉, average lateral area, 〈*A*〉, and cellular volume fluctuations, 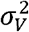, in *in vivo* tubes as a function of the apico-basal coordinate, *s* (*n* = 20 glands, similar to **Fig. 1D-E, G). D-E:** Cross-correlation analysis between energy and intercalation profiles (same as **Fig. 1H-I).** The error band indicates in this case the variability among experimental samples (*n* = 20 glands).

We determined the average basal surface ratio (thickness) of the salivary glands, 〈*s_b_*〉 = 8.5 ± 1.1, the average percentage of scutoids, 72 ± 12%, the average 3D connectivity, 〈*n*_3*D*_(*s_b_*)〉 = 6.6 ± 0.2, and the average number of apico-basal intercalations per cell, 〈*i*(*s_b_*)〉 = 1.2 + 0.3, thus confirming *in vivo* the validity of the formula 〈*n*_3*D*_〉 = 6 + 〈*i*〉/2 **(STAR Methods** and **Fig. S1).** We also calculated the spreading coefficient of the 3D connectivity, *η*^2^ = 1.2 · 10^-2^, **(Fig. 3B),** and the 2D polygonal distributions in the apical and basal surfaces **(Fig. S2).** Interestingly, we observed a small, but significant, increase of the number of hexagons on the basal surface of the wt glands (see **Fig. S2, Table S1).**

Further, in order to derive how energy contributions change as a function of the apico-basal coordinate, *s*, we implemented an algorithm that obtains the concentric radial sections of *in vivo* tubes from apical to basal (Yang et al., 2019) **(STAR Methods).** These sections were used to quantify as a function of the surface ratio, *s,* the number of apico-basal intercalations, 〈*i*(*s*)〉, the average lateral area, 〈*A*(*s*)〉, and the cellular volume fluctuations, 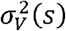 **(Fig. 3C).** Similarly to the procedure that we implemented in the Voronoi tubular model **(Fig. 1H, I)**, we used these *in vivo* data to perform a cross-correlation analysis between 〈*i*〉 and the energy contributions 〈*E_Z_*〉 (*Z* being *A* or *V,* i.e., surface tension or elastic terms). The results indicated that *in vivo* intercalations also correlate stronger with surface tension energy contributions than with elastic terms: ~98% vs. ~90% at zero (optimal) lag **(Fig. 3D).** We also found that in this case, by including the extra contribution from the apical contractile energy to the surface tension energy, i.e. 〈*E_A_*〉 + 〈*E_L_*〉), slightly decreases the correlation down to ~95% (optimal lag) but does not modify that of the elastic term, i.e. 〈*E_A_*〉 + 〈*E_L_*〉) **(Fig. 3D).** As for the cross-correlation between *∂_s_*〈*i*〉 and *∂_s_*〈*E_Z_*〉, we also found that in *in vivo* tubes it is more significant for the case of the surface tension energy, ~80%, than for the elastic contribution, ~70%. In addition, we also observed a positive lag for *∂_s_*〈*E_V_*〉 that suggests an instructive role of elastic energy variations towards changes in the number of apico-basal intercalations **(Fig. 3E).**

Subsequently, we sampled the Voronoi tubular model in terms of the disorder configuration (CVT scale) and the value of the thickness, *s_b_,* that leads to a tube that represents the aforementioned properties observed *in vivo. We* found that the V8 *in silico* model with *s_b_* = 1.75 displayed a scutoidal prevalence, 79 ± 5%, average number of 3D neighbors, 6.72 + 0.08, average number of apico-basal intercalations per cell, 1.4 + 0.1, and value of the 3D histogram spreading, *η*^2^ = 1.4 · 10^-2^, comparable to those found in *in vivo* tubes **(Fig. S3).** Further, the 2D polygonal distributions in the apical and basal surfaces of the V8 model (*s_b_ =* 1.75) were found to be similar and in agreement with those found in wt salivary glands **(Fig. S2** and **Table S1).** We also observed that the increment apico-basal transitions by means of a larger surface ratio (*s_b_* = 10) leads to an increase of topological disorder (larger variance of cell sidedness, see **Fig. S2** and **Table S1);** a phenomenon that is similar to that observed in T1-transitions (Blankenship et al., 2006; Zallen and Zallen, 2004). Finally, we implemented the cross-correlation analyses between intercalations and energy contributions in the V8 model (*s_b_* = 1.75). We obtained similar features as those obtained *in vivo,* including the suggested instructive role of elastic energy variations towards changes in the number of apico-basal intercalations **(Fig. S3).** Altogether, we concluded that the V8 model (*s_b_* = 1.75) reproduces the 2D and 3D packing properties of the *Drosophila*’s larval salivary glands.

### A biophysical model explains the cellular connectivity observed in *in silico* and *in vivo* tubular epithelia

In order to explain how the number of 3D neighbors of a cell (i.e., the cellular connectivity) changes as a function of the apico-basal coordinate, we developed a biophysical model. The model is based on a Kolmogorov rate equation and accounts for the probability of cells to increase their 3D connectivity as the radial coordinate along the apico-basal axis changes from *s* to *s* + *ds* **(Fig. 4A, B** and **STAR Methods):**

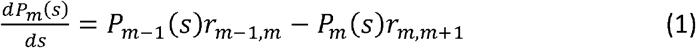

**Figure 4.**
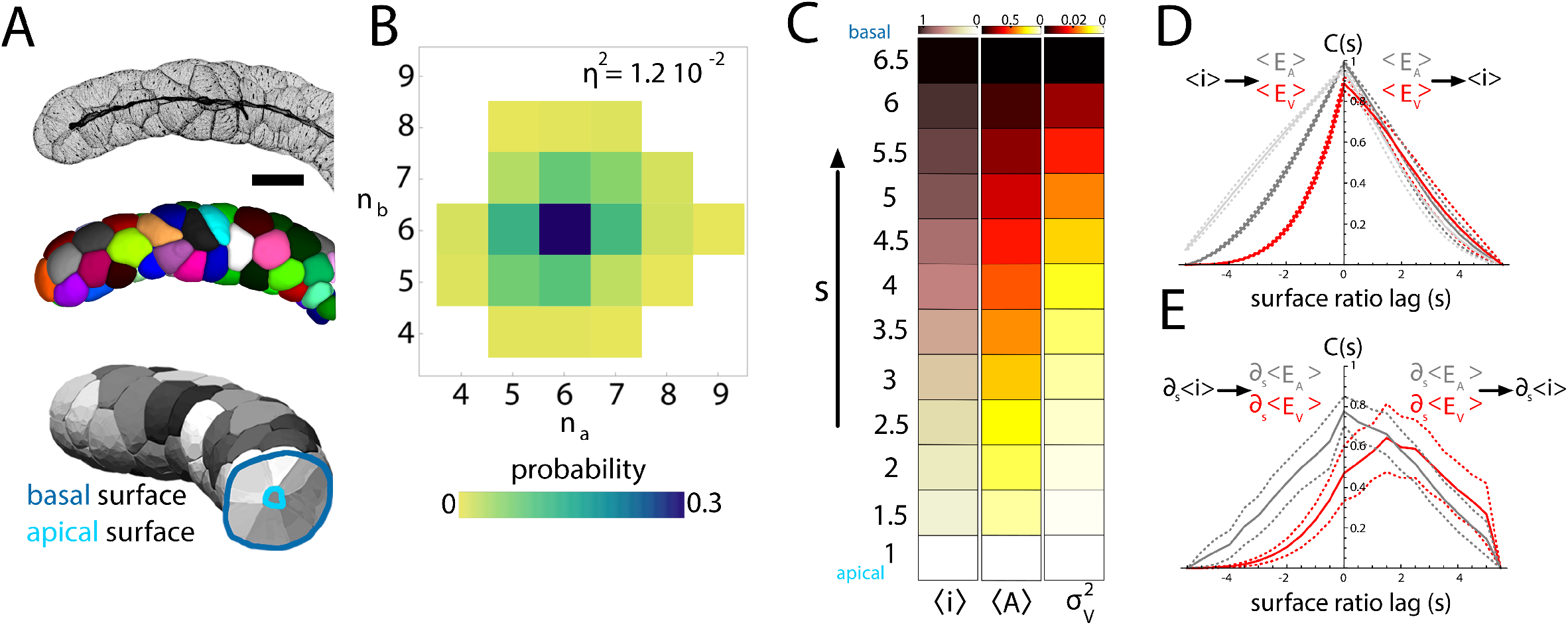
Biophysical model: *in silico* and *in vivo* results. **A:** (top) Cell intercalations along the apico-basal axis can be visualized as spatial T1 transitions (non-reversible: once a neighbor is won it cannot be lost). (bottom) The “poor get richer” principle **(STAR Methods)** suggests an increasing energetic cost (i.e., a larger activation energy) for recruiting new neighbors as a function of the number of neighbors already won. In our model, *β*(*s*) accounts for the energetic cost per 3D neighbor (per apico-basal intercalation) to recruit a new neighbor **(STAR Methods). B:** The energy landscape shown in **B** (bottom) can be modeled by a stochastic dynamics (a Kolmogorov rate equation) where cells increase their 3D neighbors with a probability per unit of surface ratio, *r_n,m_,* that depends on the activation energy, *β*(*s*), a ‘bare’ transition rate, *α*, and the maximum cell connectivity *N_max_* **(STAR Methods). C:** Results of the Kolmogorov model in V8 (*s_b_* = 1.75) *in silico* tubes (*n* = 20). The left/center density plots represent the cellular connectivity distribution (i.e., the fraction/probability of cells with a given number of 3D neighbors) as a function of *s* obtained in the Voronoi simulation (left) and as predicted by the Kolmogorov model (center); the red circles (left/right) indicate the average number of 3D neighbors per cell 〈*n*_3*D*_〉; the red line (center/right) shows 〈*n*_3*D*_〉 as obtained by the Kolmogorov model. The density plot on the right shows the difference between the predicted and the actual connectivity distributions and the corresponding error, *ε*^2^ (magenta lines). **D:** Same as panel **C** but results obtained in salivary glands (*n* = 20 glands). The maximum value of *s* in the analyzed radial sections of the glands is *s* = 6.5. This value being the largest radial section of the smallest gland **(STAR Methods).**

This equation determines, as a function of *s,* the set of probabilities {*P_m_*(*s*)} =*P*_3_(*s*), *P*_4_(*s*),⋯,*P_N_max__*(*s*), i.e., the fractions of cells with a given number, *m,* of 3D neighbors such that ∑_*m*_*P_m_*(*s*) = 1. Thus, the average 3D cellular connectivity (i.e., the average number of 3D neighbors per cell) as a function of s reads 〈*n*_3*D*_(*s*)〉 = 〈*m*〉 = ∑_*m*_*m P_m_*(*s*).

In Eq. (1), *r*_*m,m*+1_ accounts for the transition “rate” at which 3D are gained, i.e., the probability per unit of *s* to increase the cellular connectivity by one cell. By drawing parallels between apico-basal intercalations and planar T1 transitions (Bi et al., 2014; Gómez-Gálvez et al., 2018; Sanchez-Corrales et al., 2018) we assumed, following the Eyring model (Eyring, 1935), that cells need to overcome an energy barrier, Δ*E_m_*(*s*), to gain a new 3D neighbor, that is, *r*_*m,m*+1_~*e*^−Δ*E_m_*(*s*)^ (**Fig. 4A, B**). Our experimental, computational, and analytical results (see **STAR Methods** and **Figs. S4, S5)** support the idea that *r*_*m,m*+1_ = *α*(*N_max_* – *m*)*e*^-*mβ*(*s*)^ where *α* is a ‘bare’ transition “rate”, *β*(*s*) accounts for the energy cost required to gain one 3D neighbor at a given position of the apico-basal coordinate, *s,* and *N_max_* is the maximum possible 3D cellular connectivity for a cell (i.e., if *m* = *N_max_* then *r*_*m,m*+1_ = 0) **(STAR Methods).** This model predicts a logistic-like growth of the cellular connectivity **(STAR Methods).** In order to assess the validity of our model we implemented a fitting/optimization procedure that provides the value of the model parameters that minimize the error in the fitting of the curve 〈*n*_3*D*_(*s*)〉 **(STAR Methods).** Our results show an excellent agreement for all values of the CVT scale **(Fig. S6, Table S1),** the computational tubular model that represents the best in vivo data, i.e. V8 (*s_b_* = 1.75) **(Fig. 4C),** and the wt salivary glands **(Fig. 4D).** We further assessed the goodness of the biophysical model by predicting accurately the 3D neighbor’s distribution as a function of the apico-basal coordinate, {*P_m_*(*s*)} **(Fig. 4C, D, Fig. S6).**

### Genetic perturbations modify the 3D cellular connectivity properties of epithelial tubes

Our analyses suggest that surface tension is the energy contribution that affects the propensity of apico-basal intercalations the most. Surface tension energy originates in adhesion-mediated interactions between cells that ultimately modulate the magnitude of cell-cell contacts. Following these ideas, we explored the role of cell-cell adhesion by experimentally reducing the amount of the E-cadherin (E-cadh). For this aim, we overexpressed a *UAS-RNAi* line specific for the *shotgun* (*shg*) gene on the developing *Drosophila* salivary gland (Brand and Perrimon, 1993; Hammond et al., 2000; Tepass et al., 1996) **(STAR Methods).** We compared the E-cadh antibody fluorescence profiles in wt and the mutant glands (ΔEcad) and confirmed the reduction of E-cadh levels in the latter **(Fig. S7).** The cells in the ΔEcad glands bulged at the basal surface and were smaller than the wt cells **(Fig. 5A, Table S1, Fig. S7).** We processed these glands to extract their 3D cellular connectivity features and average energy profiles **(STAR Methods** and **Table S1).** We determined the effective average basal surface ratio **(STAR Methods)** of the mutant salivary glands 〈*s_b_*〉 = 7.4 ± 0.8, the average percentage of scutoids, 65 ± 14%, the average 3D connectivity, 6.5 ± 0.2, and the average number of apico-basal intercalations per cell, 1.1 ± 0.4, **(Table S1, Fig. S7).** These values confirm the validity of the formula 〈*n*_3*D*_〉 = 6 + 〈*i*〉/2 in this genetic background too. Also, the cross-correlation analysis revealed that the surface tension energy remained as the main energy contribution **(Fig. S8).** Thus, ΔEcad and wt glands reached the same 3D connectivity although the effective surface ratio of the former was smaller **(Table S1).** These results suggest that a decrease in the cellular adhesion facilitates the emergence of apico-basal intercalations.

**Figure 5.**
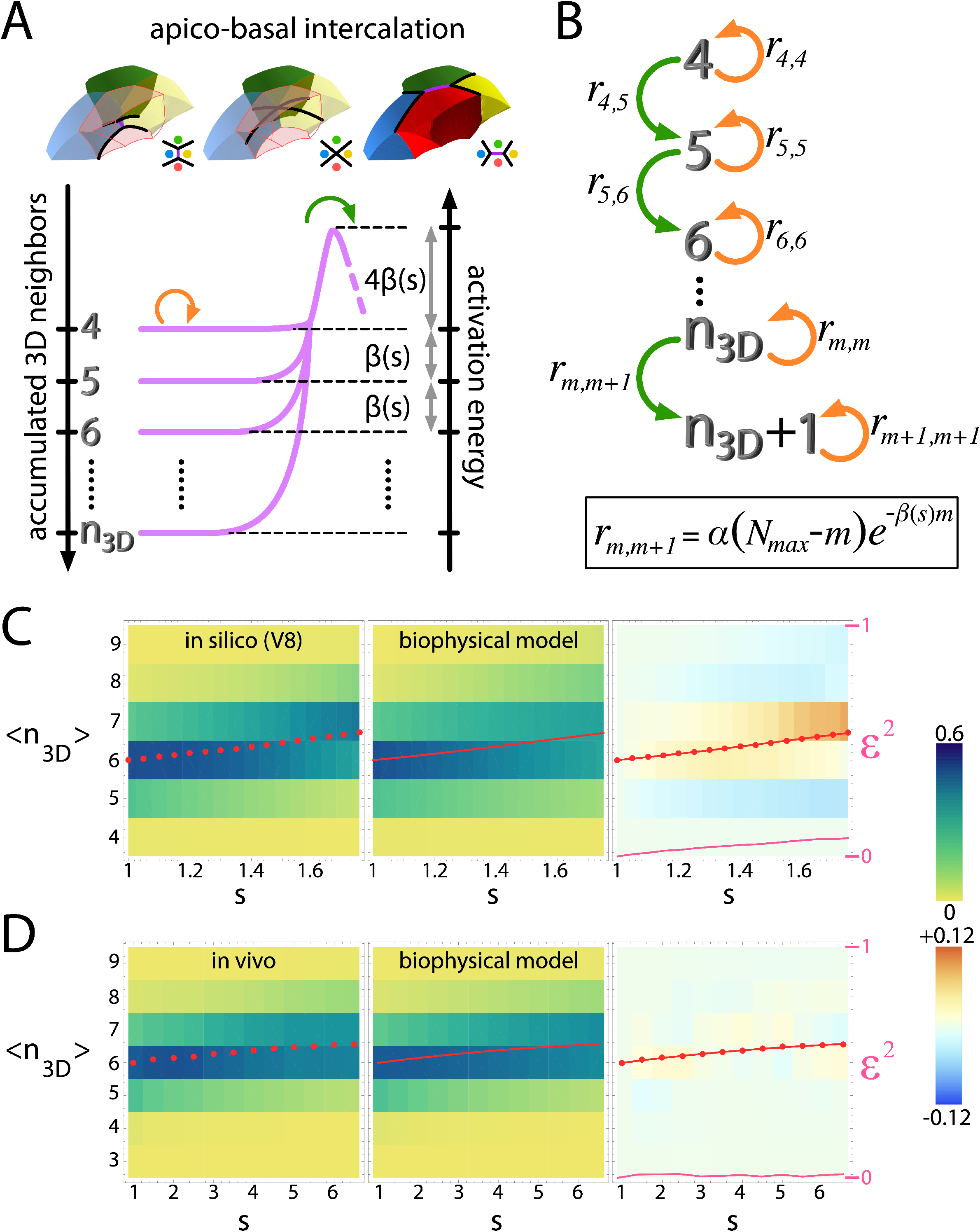
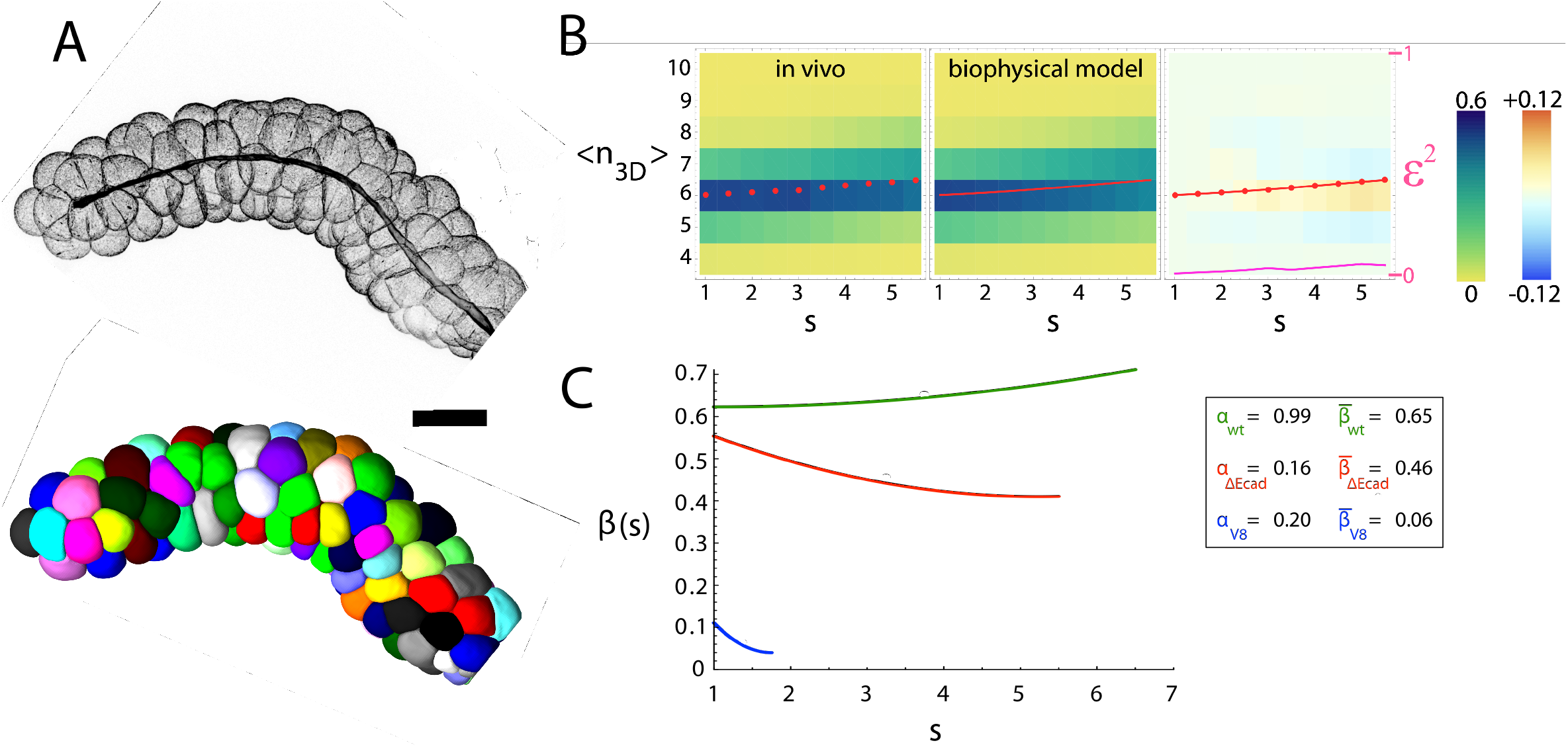
Biophysical analysis of ΔEcad salivary glands. **A:** (top) Full projection of a ΔEcad salivary gland (cell contours stained by Cy3-labeled phalloidin, **STAR Methods)**. (bottom) Computer rendering of the mutant salivary gland shown on top. Scale bar 100*μ*m. **B:** Results of the Kolmogorov model for the ΔEcad salivary gland (*n* = 10 glands). The left/center density plots represent the connectivity distribution (i.e., the fraction/probability of cells with a given number of 3D neighbors) as a function of *s* obtained in the Voronoi simulation (left) and as predicted by the Kolmogorov model (center); the red circles (left/right) indicate the average number of 3D neighbors per cell 〈*n*_3*D*_〉; the red line (center/right) shows 〈*n*_3*D*_〉 as obtained by the Kolmogorov model. The density plot on the right shows the difference between the predicted and the actual connectivity distributions and the corresponding error, *ε*^2^ (magenta lines). Same color code than in **Fig 4C-D.** The maximum value of *s* in the analyzed radial sections of the glands is *s* = 5.5. This value being the largest effective radial section of the smallest gland **(STAR Methods). C:** Energy cost required to gain additional neighbors as a function of *s* **(STAR Methods)** for wt glands (green), for ΔEcad glands (red), and for the V8 (*s_b_* = 1.75) model (blue). The inset shows the values of the bare transition rates, *α*, and the average value (along the apico-basal coordinate) of *β.*

### The reduction of cellular adhesion decreases the activation energy required to produce apico-basal intercalations

The fitting/optimization procedure of the mutant data showed, as in the case of the wt phenotype and the *in silico* tubes, an excellent agreement **(Fig. 5B)** that allowed us to estimate the energy-related parameters as summarized by *α* and *β*(*s*) **(Fig. 5C** and **Table S1).** The estimation of *α* and *β*(*s*) in wt and mutant tubes indicated that the energy required to gain an additional neighbor, *β*(*s*), is larger in *in vivo* tubes than in the computational model independently of the CVT scale **(Fig. 5C, Fig. S9, Table S1),** see **Discussion.** Finally, the results obtained from the analysis of the mutant glands confirmed that a decrease in the cellular adhesion facilitates the emergence of apico-basal intercalations since the activation energy gets reduced in the ΔEcad phenotype when compared to the wt case **(Fig. 5B** and **5C,** and **Table S1).** In particular, in the mutant case, the curve that describes the energetic cost to gain new neighbors as a function of the apico-basal coordinate, *β*(*s*), lies below the curve of the wt background **(Fig. 5C)** and we found that the average energy cost, 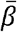 **(STAR Methods),** is ~43% larger in the wt case: 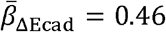 and 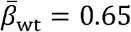. Altogether, when we challenged the biophysical model with the perturbation experiment, we obtained the expected results: smaller cellular adhesion leads to a smaller energetic cost to gain new neighbors.

## DISCUSSION

Here we have shown how a biophysical principle underlies the emergence of functionally complex 3D developmental structures. Namely, cells increase their 3D connectivity in a logistic-like fashion by means of apico-basal intercalations that require overcoming an energy barrier that grows with the number of 3D neighbors. Thus, our analyses explain how the presence of the novel paradigm of epithelial cells’ shape and packing, the *scutoid,* affects the cellular connectivity in the third dimension. In that regard, we have shown how the 3D cellular connectivity and tissue energetics are coupled, and we have proposed a quantitative biophysical model to explain that relationship.

Our biophysical model relies on a phenomenon observed in the Voronoi computational simulations, supported by mathematical arguments, and confirmed in experiments: the “poor get richer” principle (see **STAR Methods).** Roughly speaking, we have shown that the fewer neighbors a cell has on a surface, the larger is the probability of a 3D cellular connectivity increase. Interestingly, a similar idea has been reported in T1 dynamical processes during the remodeling of planar epithelia where it has been shown that the energy barrier associated with cellular remodeling, rather than being constant, depends on the cellular environment (Bi et al., 2014). Since the scutoidal geometry can be related to planar T1 transitions by exchanging the concepts of space and time, this result reinforces the idea of the existence of universal principles driving the organization of tissues.

From the viewpoint of the tissue energetics, both the Voronoi model and *in vivo* tubes, identify the surface tension energy as the main cause of scutoids appearance and hints at the elastic energy as an additional driver for modulating the propensity of cells to undergo apico-basal intercalations. In that context, our results suggest that the so-called ‘bare’ transition rate, *α*, or even the energy cost required to gain additional neighbors, *β*(*s*), would depend on both contributions. Related to this, previous studies about the role of fluctuations in the remodeling of cellular aggregates have shown that elastic behaviors (opposed to plastic ones) contribute to reduce the cell stress by lowering the energy barrier that cells need to jump over during cellular rearrangements (Marmottant et al., 2009).

In our study we have found that in real tissues (both wt and mutant) the value of *β*(*s*) is larger than in Voronoi models. We hypothesize that it is due to the purely geometrical description used in the latter. In the *in silico* model the apico-basal intercalations develop as a result of a topological constraint (a Voronoi tessellation) that we have shown describes appropriately the geometrical and packing properties of tubular epithelia. However, in the salivary glands, on top of that constraint, the cells must actively remodel their membranes and cytoskeleton to make the transitions possible. In that context, the cytoskeleton, adhesion molecules, and cellular membranes are responsible for the biophysical properties of epithelia including their energetics (Gómez-Gálvez et al., 2021b). Thus, to challenge the proposed biophysical model we measured the value of *β*(*s*) in salivary glands where the amount of the adhesion molecule E-cadh was reduced. Since the 3D connectivity necessarily increases with the surface ratio, the lower effective surface ratio of the mutant gland should correspond with a reduction of the 3D connectivity. However, our results show that wt and mutant glands present the same value of 〈*n*_3*D*_(*s_b_*)〉 **(Table S1, Fig. S7),** thus indicating that the decrease of adhesion facilitates the appearance of apico-basal intercalations. In terms of the tissue energetics, these results suggest a reduction of the energy barrier required to undergo apico-basal intercalations in the mutant glands. This prediction was confirmed by the biophysical model that provided a lower *β*(*s*) in the mutant case compared with the wt.

As for the technical advances associated to our work, we point out that a high level of detail is necessary to quantify the apico-basal intercalation phenomenon and to compare the *in vivo* data with computational models (Gómez-Gálvez et al., 2021a). Along these lines, the importance of a realistic analysis of 3D cell-cell contacts has been highlighted by recent studies focused on understanding the growth of mouse embryonic lung explants (Gómez et al., 2021) and the early development of *C. elegans* (Cao et al., 2020) and Ascidians (Guignard et al., 2020). Our novel methodological pipeline **(STAR Methods)** allows to implement the accurate 3D reconstruction of cells in epithelia subjected to curvature. This analysis makes possible to quantify how different packing properties, e.g., intercalations, depend on the apico-basal coordinate. These technical improvements are necessary to extract biological consequences about the cellular and mechanical basis of self-organization in curved tissues (Ambrosini et al., 2017; Hirashima and Adachi, 2019; Inoue et al., 2019) or even whole embryos (Shahbazi et al., 2019).

As for the broader implications of our findings, our results provide new biological insight into the regulation of cell-cell connectivity in curved tissues. This property ultimately regulates juxtracrine signaling, and is pivotal for early development, primordia patterning, and cell fate determination (Guignard et al., 2020; Sharma et al., 2019; Tung et al., 2012). Moreover, recent research has shown that cellular connectivity regulates the viscosity of tissues (Petridou et al., 2021).

Therefore, our findings open new ways to draw implications about primary developmental processes in which epithelial bending is essential such as tubulogenesis, gastrulation, neurulation, and early embryo development. In addition, we argue that, while our analyses focus on static tissues, our results could also be relevant to understand active 3D tissue remodeling. Dynamic changes of *β*(*s*) would modify the apico-basal intercalation propensity and therefore the material-like properties: the larger *β*(*s*) the more solid-like the tissue would behave.

Recent studies have confirmed that adhesion-dependent active remodeling can be connected to an increased activity of neighbor exchanges. In particular, loss of function mutants of N-cadherin in the presomitic mesoderm of the zebrafish embryo cause an increase in extracellular spaces and a solid-fluid jamming transition (Mongera et al., 2018). In addition, it has been recently shown that the stabilization of E-cad at the cellular junctions in the *Drosophila eye* drives an increase of tension that can be transmitted across the tissue (Founounou et al., 2021). This tension results in a reinforcement of the solid-like tissue behavior. The salivary glands experiments confirm that a reduction of E-cadherin increases the apico-basal intercalation propensity. Our biophysical model predicted that such an increase of the apico-basal intercalation propensity must be correlated with a decrease of the energy barrier *β*(*s*). Notably, this prediction was confirmed through the biophysical analyses of the ΔEcad samples.

Finally, with respect to the applicability of our results to other areas, we expect that the emerging field of organoids will benefit from our discoveries. A precise quantification of the 3D connectivity could then help to understand the lack of reproducibility in organoid production, one of the biggest challenges of the field (Clevers, 2016; Huch et al., 2017; Schutgens et al., 2019). Also, from a medical point of view, it has been recently shown that tissue curvature affects tumor progression due to the imbalance of tensions in apical and basal surfaces of epithelial tubes (Messal et al., 2019). Our study explains how cell energetics affect the 3D packing of these cells and therefore may shed light on the mechanism of tumorigenic morphogenesis in tubular organs.

## ACKNOWLEDGMENTS

L.M.E., P.V.-M. and P.G.-G. has been supported by the Ramón y Cajal program (PI13/01347); L.M.E., P.V.-M. and P.G.-G. work is funded by the Ministry of Economy, Industry and Competitiveness grant BFU2016-74975-P co-funded by FEDER funds, and by the Spanish Ministry of Science and Innovation Ministry of Science through grant PID2019-103900GB-I00. A.T. is funded by a “Contrato predoctoral PIF” from Universidad de Sevilla. L.M.E. and J.A.A.-S. work is funded by the Junta de Andalucía (Consejería de economía, conocimiento, empresas y Universidad) grant PY18-631 co-funded by FEDER funds. I.A-C. would like to acknowledge the support of the 2020 Leonardo Grant for Researchers and Cultural Creators, BBVA Foundation, and also the University of the Basque Country UPV/EHU grant GIU19/027. S.A. and J.B. have been supported by a Faculty Innovation Grant (FIG) by Lehigh University JB-FIG-2019. J.B. also acknowledges financial support from the Spanish Ministry of Science and Innovation through grant PID2019-105566GB-I00 and from the *LifeHUB* Research Network (CSIC).

## DECLARATION OF INTERESTS

The authors declare no competing interests.

#### Box

**A: Scutoids** are prismatic-like geometric solids bounded between two surfaces (top and bottom). Scutoids are characterized by three main properties: i) The shape of their top and bottom bases, and of every parallel section between them, are polygons. ii) The lateral surfaces of scutoids can be concave and/or convex surfaces such that a set of scutoids can be packed together (laterally) without leaving any empty space. iii) Scutoids have at least one vertex along the top-bottom axis such that when packed together there are changes in the nearest-neighbors relationship. The example shows a scheme of a stereotypical scutoid (left) and four scutoids packed together (right).

**B:** A **T1-transition** is a tissue rearrangement observed in epithelial surfaces where a 4-cells’ motif swaps nearest neighbors along time. T1-transitions enable tissue plasticity through cellular reorganizations that lead, for example, to elongation in developing tissues.

**C:** An **apico-basal transition,** aka an **apico-basal intercalation,** is a tissue rearrangement along the apico-basal (top-bottom) axis of cells that lead to new cellular contacts (nearest-neighbors exchange). An apico-basal transition is similar to the T1-transition but instead of developing along the time it does along space.

**D:** A 2D **tessellation** (aka a 2D mosaic) is a partition of a surface with tiles that do not overlap or leave any gaps. In this example, the tiles are octagons and squares.

**E:** A 2D **Voronoi diagram** is a particular type of tessellation built by convex polygons (Voronoi cells). These polygons emerge from a set of generator seeds (black points), such that each cell contains the region that is closer to its generating seed. The so-called **Lloyd’s algorithm** makes the seeds of a Voronoi diagram to converge progressively to the centroids (blue points): once a Voronoi diagram is obtained for a set of seeds (black Voronoi diagram, *V_N_*), an iteration of the Lloyd’s algorithm consists in repeating the Voronoi tessellation by replacing the seeds by the centroids of the Voronoi tiles (blue Voronoi diagram, *V*_*N*+1_). The Lloyd’s algorithm makes the Voronoi diagram progressively more ordered in terms of the polygonal distribution of the Voronoi cells.

**F:** The **surface tension energy** is related to the cell-cell adherence through their lateral area contacts. For each cell, the surface tension energy reads, *E_A_* = Λ*A*, where Λ and *A* are the effective surface tension parameter and the cellular lateral area respectively. Thus, the average surface tension energy of cells reads 〈*E_A_*〉 = Λ〈*A*〉 = *E*_〈*A*〉_ and is independent of the fluctuations of *A*.

**G:** The **contractile energy** is related to the polarized cortex activity of epithelia cells at the apical surface. The contractile energy reads *E_L_* = Γ*L*^2^, where *L* stands for the cellular perimeter at the apical surface and Γ is the cortical tension energy per unit of cell apical area. As a result, the average cell apical contractile energy increases with the fluctuations of the apical perimeter: 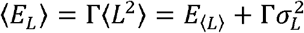 where 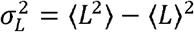 is the cellular apical perimeter variance.

**H:** The cell **elastic energy** is related to the volume conservation of cells. The cell elastic energy reads, 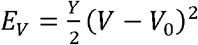, where *Y* is proportional to the Young modulus (a quantification of the relationship between the cellular stress and strain) and, *V* and *V*_0_ represent the *actual* and target cellular volumes respectively. The average elastic energy per cell increases with the fluctuations of the volume: 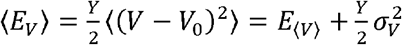 where 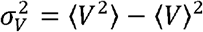 is the cellular volume variance.

## STAR METHODS

### Immunohistochemistry and confocal imaging of salivary glands

Flies were grown at 25 °C using standard culture techniques. The following lines were used: *Oregon R (wt), UAS-shg-RNAi* (BL38207), *AB1-Gal4* (BL1824). *AB1-Gal4* drives Gal4 protein in the third instar larva salivary gland. We dissected the salivary glands from third instar larvae. After PBS dissection, the glands were fixed using 4% paraformaldehyde in PBS for 20 min. The samples were washed three times for 10 min with PBT (PBS, 0.3% Triton) and then incubated for 1 hr 45 minutes at room temperature with Cy3-labeled phalloidin (Sigma) to label the cell contours of the epithelial cells. Stained larval salivary glands were mounted using Fluoromount-G (Southern Biotech). We used two pieces of double-sided adhesive tape (one on top of each other) as a spacer (Aldaz et al., 2013), so the salivary glands preserve their shape. Images were taken using a Nikon Eclipse Ti-E laser scanning confocal microscope. The images were captured using a ×20 dry objective and 2.5 μm steps between slices. The image stacks were exported as 1024 × 1024 pixels TIFF files.

### Quantification of fluorescence intensity

The E-cadherin fluorescence intensity was measured in Fiji by using the Plot Profile tool. We used 3 wt and 3 ΔEcad representative glands, taking 10 individual measurements for each sample **(Fig. S7).** We used rectangular ROIs to measure the intensity profiles of lateral cell membrane in the Z-depth where the lumen was visible. In this way, we were able to capture the whole lateral cell membranes from apical to basal. To ensure, a high-quality detection of the cell membrane we developed a maximum Z-projection of those Z-slices where the cell outline of interest and the lumen are clearly visible. Note that the output of the Plot Profile is a 2D plot that displays a “column average plot”, where the *X* axis represents the horizontal distance through the selection (apico-basal cell outline) and the *Y* axis the vertically averaged pixel intensity.

### 3D glands segmentation

To segment the salivary gland stacks of images and reconstruct (semi-automatically) the shape of cells in three dimensions we used the Fiji (Schindelin et al., 2012) plugin LimeSeg (Machado et al., 2019). It infers cell outlines by using surface elements (“Surfels”) obtained by manually placing single ellipsoidal-like seeds on every cell (see https://imagej.net/LimeSeg for details). Once cell outlines were found, we exported them as point clouds (output). We developed a custom-made Matlab code (2021a MathWorks) to postprocess the output of LimeSeg in order to correct errors and obtain perfectly segmented salivary glands. In addition, we manually segmented the lumen of the glands using the Volume Segmenter app, in Matlab. To faithfully represent the gland as a cylinder, we selected a subset of cells: cells that were not ductal, neither located at the tip of the gland.

To segment mutant salivary glands we took advantage from the 20 segmented wt salivary glands, and we used them as training dataset into a deep-learning segmentation pipeline. We trained a stable 3D-U-Net CNN ((Franco-Barranco et al., 2021), https://github.com/danifranco/EM_Image_Segmentation) using as input the salivary glands phalloidin channel (actin filaments) staining cell outlines, and as target the segmented cell outlines. The output (prediction) of this pipeline was a probability map of cell outlines, that was post-process using the PlantSeg (Wolny et al., 2020) segmentation module to extract individual instances. Here, again, we segmented the lumen of the glands using the Volume Segmenter app, and segmentation errors were curated using our custom-made Matlab code.

To obtain the cellular neighborhood relations of salivary glands for different values of the radial expansion, we proceeded as follows. We calculated the cell height by estimating the distance between the centroid of the cell apical surface with respect to the centroid position of its basal surface, *d*(*s_a_, s_b_*). Then, to capture a concentric radial section of the gland, we linearly extrapolated the equivalent cell height to the given surface ratio, *s*:

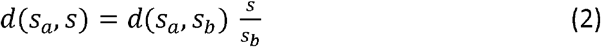

where *d*(*s_a_,s*) is the Euclidean distance between the position of the centroid of the cell at the apical surface, *s_a_* = 1, and the position of the centroid at a value *s* = *R/R_a_* of the radial expansion. Finally, to obtain the gland cylindrical radial section for a given value of the radial expansion, *s*, we collected all voxels between apical and the upper bound of the calculated distance *d* (*s_a_, s*).

### Salivary glands measurements

We quantified the following geometrical and topological/connectivity descriptors of the segmented salivary glands using a custom-made Matlab code:

- Surface ratio (*s*): Assuming a cylindrical shape for glands, we estimated *s* by measuring the minimum distance between each cell apical centroid and lumen skeleton (*R_a_*), and measuring *h,* the distance between apical cell centroid and cell centroid of an outer cell layer (i.e., basal surface or a concentric layer between apical and basal). Being, the individual surface ratio of a cell, 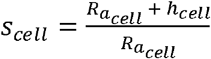, we averaged all the individual cell measurements to calculate the representative *s* value corresponding to a gland, *s* = 〈*s_cell_*〉.
- Cell apical perimeter, lateral surface area, and the cellular volume.
- Number of cell contacts: we measured the number of cell neighbors of each cell surface, that is, apical, basal or lateral. In order to remove artefacts, 2 cells must share at least 0.5% of their lateral surfaces area to enable them to be considered as neighbors. *n_a_, n_b_,* and *n*_3*D*_ of each gland were calculated after averaging the number of cells neighbors along the gland.
- Percentage of scutoids, average of apico-basal transitions. We quantify the percentage of scutoidal cells that conform the gland and the number of apico-basal transitions in which each cell is involved.

All the measurements were carried along different concentric radial sections of the glands. We captured the gland thicknesses starting at the apical surface (*s_a_* = 1) and increasing progressively the surface ratio by Δ*s* = 0.5, until reaching the basal surface (*s_b_*). In this way, the number of the captured radial sections will depend on the *s_b_* value of each gland. To compare the glands of each phenotype (either wt or mutant) in terms of connectivity related with the surface ratio, we used a maximum radial section common to all the glands.

In ΔEcad mutant glands, due to their phenotype (cells bulge at the basal surface), we removed the bulging tips of cells to quantify the *effective* surface ratio *s*\*: the maximum value of the surface ratio up to which cells are contacting (that is, 1 ≤ *s* ≤ *s*\* ≤ *s_b_*). We noticed that the 3D connectivity of cells is not modified by this approach. To remove the volume of cell tips, we captured all lateral and apical surfaces of cells and we filled each cell volume using the *alphaShape* Matlab function.

### Voronoi tubular model

Using custom-made Matlab code we generated a Voronoi model that simulates the surface of a cylinder unfolded over the Cartesian plane, see details in Gomez-Galvez et al. ((Gómez-Gálvez et al., 2018), **Methods).** The only difference with the cited methodology, is that in this work the Voronoi diagrams has been constructed by means of the Delaunay triangulation technique. Therefore, we just considered the cells’ vertices information (Cartesian coordinates and connections) for a much faster computation. For each realization, we used an initial set of 200 randomly located seeds on a rectangular domain of 512 (X axis; transverse axis of cylinder) per 4096 (Y axis; longitudinal axis of cylinder). We performed this procedure for 10 different initial Voronoi diagrams (Voronoi 1 (V1, random seeds) to Voronoi 10 (V10, more ordered and homogeneous cells). These diagrams represent the apical (inner) surfaces of computational tubes, and they were obtained by applying N-1 times the Lloyd’s algorithm (Lloyd, 1982) to the random seeds, where N is then the resulting Voronoi model. For instance, to compute a V1, we use purely random seeds, while to obtain a V4 diagram, it would be required to apply 3 times the Lloyd’s algorithm to random seeds. In the limit of the CVT scale (iterations of the Lloyd’s algorithm) going to infinity the organization of cells tends to a hexagonal lattice. Subsequent radial sections that define computational tubes with different surface ratios were obtained by implementing a radial projection of the Voronoi seeds. For each apical surface of the tube, we generated 40 expansions by incrementing the surface ratios (*s_b_*) using 0.25 steps: 1 (apical), 1.25, 1.5, …, 10 (19 *s*-steps × 10 apical cell arrangements × 20 realizations).

As for the 3D reconstruction of cells in Voronoi tubes, each set of seeds that characterizes cells on a given cylindrical section defines a unique 2D Voronoi diagram at every surface and hence the corresponding 2D cellular domains. The set of 2D Voronoi regions that belong to the same radially projected seed from the apical to the basal surface then define each 3D cellular shape. Each of the obtained 3D cells was further processed using the Matlab function *‘alphaShape’* to transform the set of voxels into a compact, solid, object. This reconstruction pipeline was implemented using Matlab (2021a).

As for the connection of the CVT scale with the average elastic energy of cells, we first notice that for a given tube of length *L,* radiuses *R* and *R_a_,* and with a fixed number of cells, *N,* the average cell volume, 〈*V*〉, is independent of the CVT scale: 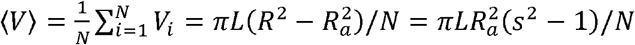. On the other hand, if cells have a target volume *V*_0_ then the elastic energy (linear regime) of cell *i* reads 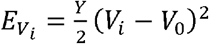, where *Y* is proportional to the Young’s modulus. Consequently, the average cell elastic energy, 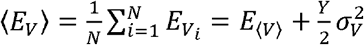, where 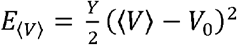 is the elastic energy of a cell with an average cell volume 〈*V*〉 and 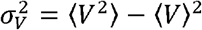 is the variance of the cellular volume (the cell size fluctuations). Since *E*_〈*V*〉_ is independent of the CVT scale and 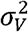 decreases with the CVT scale (i.e., as the tissue becomes more ordered) then the average elastic cell energy necessarily decreases as the CVT increases. In our simulations and experiments the cellular volume is computed by using the value of cell area sections 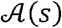 as a function of the surface ratio, *s*. Specifically, we used the trapezoidal rule, 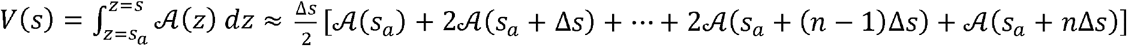. Where *s_a_* + *n*Δ*s* = *s* and Δ*s* = 0.25. Cell volumes where normalized considering Voronoi 1 tubes from CVT scale as reference, such its average cell volume will represent the unity 〈*V*(*s_b_* = 10)〉 = 1. Likewise, for estimating the surface lateral area we used the trapezoidal rule using the value of the cellular perimeter, *L*(*s*), that is: 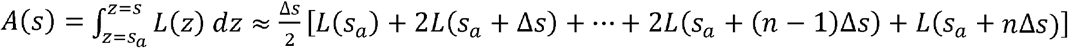. Besides, we normalized the lateral surface area following the same criterion than with volumes. Additionally, we proceed in a similar way to estimate the cellular lateral area and volume as a function of *s* in salivary glands.

### Cross-correlation definition

Dimensionless cross-correlation, *C*(*s*), between *X*(*s*) (e.g. 〈*E_A_*(*s*)〉) and *Y*(*s*) (e.g. *Y*(*s*) = 〈*i*(*s*)〉) is defined as follows: 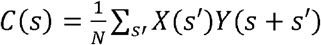 where 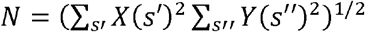 is a normalization constant such that the auto-correlation becomes one (at maximum) at zero lag. When required, spatial derivatives were estimated as 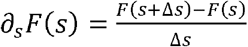.

### Voronoi tubular model measurements

We measured the following properties of cells in Voronoi tubular models: area, perimeter and number of sides of cells for a given radial section, and total number neighbors. Additionally, we computed the percentage of scutoids, the number of apico-basal transitions, the polygon distribution of every surface (radial sections). In these quantifications, we disregarded cells at the boundaries (tips of tubes) to avoid ‘border effects’.

### Relation between total accumulated 3D neighbors and the number of intercalation events

Scutoids have a Euler characteristic *χ* = 2 such that *V* – *E* + *F* = 2, where *V, E,* and *F* accounts for the number of vertexes, edges, and faces respectively. We assumed that the apical, *a*, and basal, *b,* faces of scutoids tessellating a cylindrical space have radial coordinates *R_a_* and *R_b_* respectively. Then, for any value of the surface ratio expansion, *s* = *R/R_a_*, these solids can be mapped into a connected plane graph with the same Euler characteristic (a sort of projection of the vertexes and connectors into the plane, see **Fig. S10.** Thus, as a function of *s*, the accumulated number of 3D neighbors reads *n*_3*D*_(*s*) = *E*(*s*) – *V*(*s*). Since in tubular geometries the radially projected seeds from the apical to the basal surface never come closer, as s increases (i.e., apico-basal intercalations are not reversible).

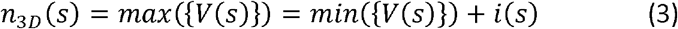

where {*V*(*s*)} = {*V*(1), *V*(1 + *ds*),⋯,*V*(*s_b_*)ߛ and *i*(*s*) denotes the number of intercalation points in the interval *s* ∈ [1,*s_b_*]. In the case of a 3D tessellation with *N* cells, where *M* of them do not show any intercalation, the total number of accumulated neighbors reads,

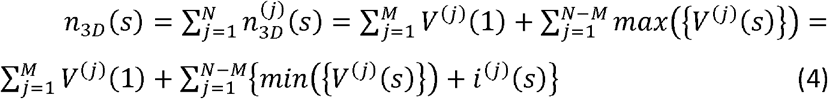

Given that each intercalation point is shared by four cells, two of them necessarily increase their number of vertices in a given *s*-plane and two of them decrease their number of vertices. Thus, in the case of a decrease *max*({*V*^(*j*)^(*s*)}) = *V*^(*j*)^(1) and in the case of an increase min({*V*^(*j*)^(*s*)}) + *i*^(*j*)^(*s*) = *V*(1) + *i*^(*j*)^(*s*). Consequently,

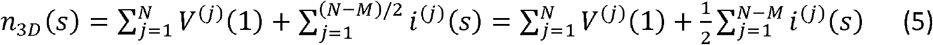

where we used the fact that for every intercalation event that increases by one the number of neighbors there is one that decreases the number of neighbors in the same amount; consequently, we can add up all intercalation events and divide by two. Hence the average number of accumulated 3D neighbors, 〈*n*_3*D*_(*s*)〉 = *n*_3*D*_(*s*)/*N* reads 〈*n*_3*D*_(*s*)) = 〈*V*(1)) + 〈*i*(*s*)〉/2; 〈*i*(*s*)〉 being the average number of apico-basal intercalations per cell. Finally, by considering that any *s*-surface, and in particular the apical surface *s* = 1, corresponds to a 2D tessellation of convex polygons, 〈*V*(1)〉 = 6 we conclude that,

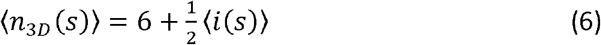

### The 3D neighbor’s accumulation in tubular epithelia follows a “poor get richer” principle

In order to investigate additional phenomena that could help to understand how the 3D cellular connectivity is regulated, we computed the net gain of cellular neighbors in epithelial tubes as a function of the 2D polygonal cell class at the apical surface. We observed, both in the salivary glands and in the Voronoi model (in particular in the case V8 (*s_b_* = 1.75) that compares the best with *in vivo* glands), that the smaller the number of neighbors of a cell at the apical surface, the larger the net gain of 3D cellular contacts **(Fig. S4).** This behavior was also obtained with respect to the 2D polygonal cell class at the basal surface **(Fig. S4).** These results suggest that, in terms of the cellular packing, tubular epithelia follow a “poor get richer” principle: the less neighbors a cell has in a surface (apical or basal), the larger the net increase of 3D cellular contacts.

### In Voronoi tubes the net gain of 3D neighbors is bounded

The “poor get richer” behavior can be justified by mathematical arguments that show that the probability to increase the cellular connectivity necessarily decreases with the number of current neighbors **(Fig. S5).** Assuming a cylindrical geometry (e.g., epithelial tubes), each point at a given radial surface can be represented into the Cartesian plane; where coordinate *x* accounts for the cylindrical transversal coordinate and coordinate *y* for the longitudinal one (see **Fig. S5).** Thus, if the coordinates of a point (e.g., a Voronoi seed) at the apical surface are given by (*x,y*), the coordinates of that point at a surface with a value of the cylindrical radial expansion *s* ∈ [1, ∞) can be found by defining the function *f_s_*: ℝ^2^ → ℝ^2^ *f_s_*(*x,y*) = (*sx,y*). Under these conditions, we aim to characterize the seeds that generate scutoids (exchanges in the neighboring relations of seeds) ass changes.

#### Lemma 1.

Given three non-colinear points {*A,B,C*} that define a circle (a nearest-neighbors relation), and another exterior point *D,* if *s* > 1 exists such that *f_s_*(*D*) is interior to the circle defined by {*f_s_*(*A*),*f_s_*(*B*),*f_s_*(*C*)}, then *D* is inside of the vertical parabola containing {*A, B, C*} **(Fig. S5).**

#### Remark.

If two of the three points {*A,B,C*} are on the same vertical line, then the parabola considered in Lemma 1 degenerates as a vertical strip. Even in this case, the thesis of the Lemma is true if we replace the interior of the parabola by the inside of the strip.

#### Proof.

Without loss of generality, we can suppose that {*A,B,C*} are counterclockwise oriented and that they have Cartesian coordinates (*a*_1_, *a*_2_), (*b*_1_, *b*_2_)and (*c*_1_, *c*_2_) respectively.

Thus, the point *D*(*x,y*) is outside the circle defined by {*A,B,C*} if, and only if, the sign of the following determinant is negative:

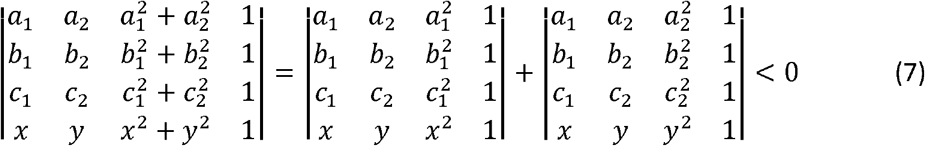

For the sake of simplicity, we represent the previous equation as:

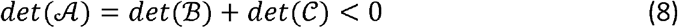

On the other hand, by considering *x* and *y* as variables, the equation 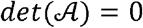 corresponds to the circle defined by {*A,B,C*}, and 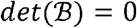 corresponds to the vertical parabola defined by the same three points. Consequently, the inequality 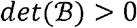 defines the locus of interior points to that parabola.

Now, assuming that *s* > 1 exists such that *f_s_*(*D*) is interior to the circle defined by {*f_s_*(*A*),*f_s_*(*B*),*f_s_*(*C*)}.Then,

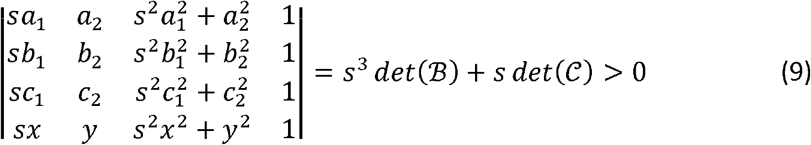

Or, equivalently, 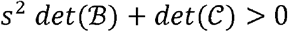, so, 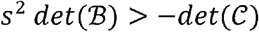. If 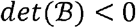, then 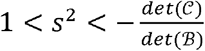 and therefore 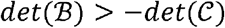. The latter is in contradiction with 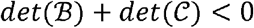. As a result, 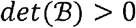, and the following inequality holds,

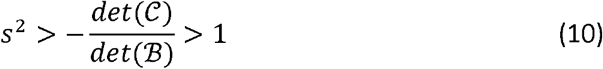

Notice that if the circle defined by {*A, B, C*} is surrounded by a set of points and we change continuously the parameter *s* in the interval [1, ∞), it is possible to detect the first point touching the circle defined by {*f_s_*(*A*), *f_s_*(*B*), *f_s_*(*C*)}. That point can be obtained by computing all the points at 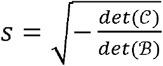. Hence, the first point contacting the circle will be that with the minimum value of *s*.

As for proving that the average of the number of neighbors of a cell induced by a seed grows is bounded as a function of the surface ratio, we state the following proposition:

#### Proposition 1.

Given a Voronoi seed representing a cell, if *n*_3*D*_(*s*) is the total number of accumulated cell neighbors as *s* increases from *s* = 1 (apical surface) to a given value of *s*, then 〈*n*_3*D*_(*s*)) is a bounded function for a finite cylinder.

#### Proof.

We model the apical surface as the cylinder 2*πr* × *h*, where *r* representes the inner radius and *h* the length of the cylinder. Given a seed *A* in that surface, in the corresponding Delaunay triangulation it appears as a point surrounded by triangles defining the neighborhood of *A.* By Lemma 1, each triangle defines a vertical parabola and a circle. So, any other seed touching *A* in other layer must be inside of one of the parabolas and outside of all circles (see **Fig. S5).** Let’s denote 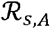 the feasible region for a new neighbor of *A* in the layer represented by *s,* i.e., all points inside one of the parabolas and outside all the circles. Thus, if 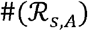 is the number of seeds in that region that are not neighbors of *A* in the apical surface, obviously, an upper bound to the number of new neighbors to 4 is given by 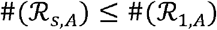.

On the other hand, that number of seeds is, in average, proportional to the density of seeds times the area of 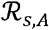, therefore, the average number of accumulated neighbors of *A,* denoted as 〈*n*_3*D*_(*A*)〉, will be bounded by the change of the density of points when growing *s,* this is to say,

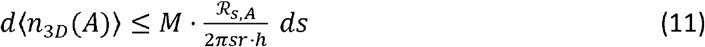

where *M* represents the total number of seeds (i.e., the total number of cells that is a constant) and the quotient is the area of 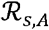 divided by the area of a given radial layer. In general, it is not possible to integrate equation (4), since the area of 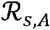 is known only in very few, particular, cases.

In the case of a finite cylinder, 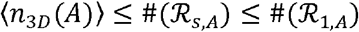 leads, summing up to all the seeds and dividing by *M,* to the upper bound

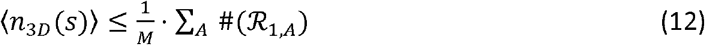

thus, 〈*n*_3*D*_(*s*)〉 is necessarily a bounded function. This expression indicates that the number of new neighbors when increasing *s* exhausts since the number of cells is a resource shared by all the layers. It is possible to obtain an upper bound to *N_max_* = lim_*s*→∞_〈*n*_3*D*_(*s*)〉 since, after a flip in the Delaunay triangulation, the edge disappearing (i.e., a cell contact loss) can never be recovered in a cylindrical geometry. Thus, *M* · (*N_max_* – *n*_3*D*_(1)) is bounded by the number of edges that complement the original Delaunay triangulation on the apical surface, that is,

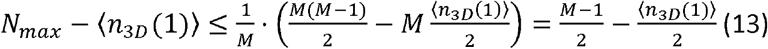

leading to

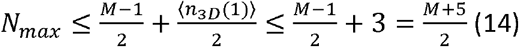

Where we have assumed that 〈*n*_3*D*_(1)〉 = 6. The simulations of the computational Voronoi model and the data of the salivary gland show that *N_max_* is in fact much smaller that the theoretical bound 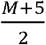.

### A Kolmogorov rate equation for the 3D cellular connectivity

The equation for how the probability, *P_m_,* of having *m* accumulated 3D neighbors (i.e., *m* = *n*_3*D*_) changes as the surface ratio (apico-basal dimensionless radial coordinate) increases from *s* to *s* + *ds* can be described by the following Markov equation **(Fig. 4A-B),**

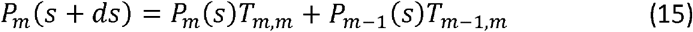

where *T_n,m_* is the probability of changing the number of neighbors from *n* to *m* due to an apico-basal intercalation. Since ∑_*m*_*T_n,m_* = 1 (normalization of the transition probabilities) and *T_n,m_* = *f*(*n,m*){*δ*_*n*–1,*m*_ + *δ*_*n,m*+1_} (each intercalation can only possibly induce to win one neighbor) then *T_m,m_* = 1 – *T*_*m,m*+1_ and the above Markov equation can be written as a Kolmogorov equation (a.k.a. Master equation):

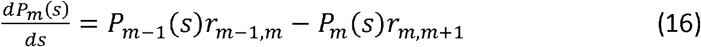

where *r_n,m_* accounts for the probability of apico-basal intercalations per unit of surface ratio, i.e., *T_n,m_* = *r_n,m_ds*. We point out that our model accounts for apico-basal intercalations that occur due to curvature effects such that *s* = *R/R_a_* changes along the apico-basal coordinate of cells (i.e., bending, folding). Thus, our model does not capture the apico-basal intercalations that develop due to active cellular processes (e.g. cellular extrusion, cell divisions) either in bended tissues or in the case of tissue planar geometries.

By following the Eyring model (Eyring, 1935), i.e., if we assume an Arrhenius-like kinetics such that to win neighbors there is an energy cost (see (Bi et al., 2014)) then 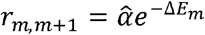, where 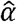 is the so-called pre-exponential factor that modulates the “bare” frequency of intercalations (per unit of surface ratio expansion, *s*) and *ΔE_m_* is a dimensionless activation energy (in units of the effective thermal energy associated with membrane fluctuations *ξ* (Marmottant et al., 2009). The observed “poor get richer” behavior suggests that the activation energy, *ΔE_m_,* increases with *m.* This can be explained as a result of a cumulative process if we assume that each neighbor that is gained implies to overcome an energy barrier, *β*(*s*), through an apico-basal intercalation. Consequently, 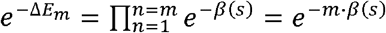. Thus, *β*(*s*) represents the dimensionless activation energy of a cell per 3D neighbor or, in the context of the different energetic contributions reviewed in this manuscript, to the energy barrier required to perform a spatial T1-transition following a surface energy minimization process (Gómez-Gálvez et al., 2018; Mughal et al., 2018). As for the dependence of *β* on *s,* the simplest mathematical form that recapitulates the fact that the apical and basal surfaces accumulate more cell-cell adherent complexes (either in wt or mutant phenotypes) is quadratic **(Fig. S7):** 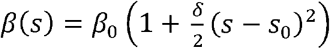. The average, along the apico-basal coordinate, of the energy cost then reads 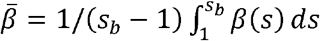. On the other hand, the mathematical calculations indicate that the intercalation rate *r*_*m,m*+1_ becomes null for a finite value of *m* or, alternatively, that the activation energy becomes infinite for a finite value of *m.* Otherwise, the net gain of new neighbors is not bounded. This fact can be accounted for by assuming that the bare frequency is a function of the number of neighbors, 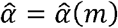, such that 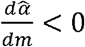 and becomes null for a finite value of *m.* For the sake of simplicity, we assume that up to first order in 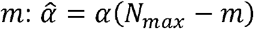, where *N_max_* is the asymptotic, maximum, number of 3D neighbors a cell can possibly have. Summarizing, the apico-basal intercalation rate *r*_*m,m*+1_ reads,

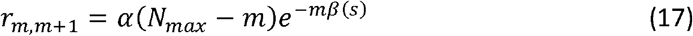

Under these conditions, the Kolmogorov equation reads,

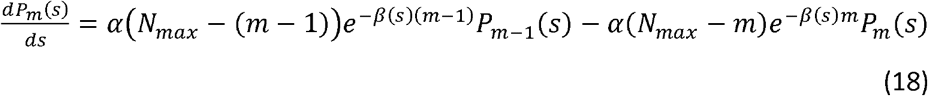

On the other hand, the equation satisfied by the average number of accumulated 3D neighbors, 〈*n*_3*D*_〉 = 〈*m*〉, reads,

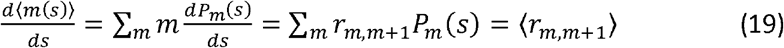

We notice that this equation implies an important role of the disorder (i.e. the distribution *P_m_*): even in conditions under which the transition rate, *r*_*m,m*+1_, is “large”, the resulting growth of 3D neighbors, 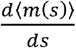 and, consequently, the net accumulation of 3D neighbors, can be more prominent in conditions where the transition rate is “small”. To illustrate this effect, we consider the following example. For the sake of simplicity, we evaluate the initial growth of 3D neighbors starting from the apical surface, i.e. we particularize Eq. (19) to the case s = 1 (and hence according to Euler’s formula 〈*P_m_*(1)) = ∑_*m*_*mP*_*m*_(1) = 6) and consider two possible conditions: a fully ordered (o) distribution with 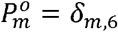 (i.e., all hexagons) and a disordered (*d*) condition that combines with equal probability cells with 3, 6, and 9 sides in the apical surface, i.e. 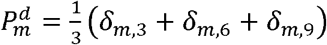. We also assume that 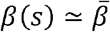 (i.e., we approximate the energy cost to gain new 3D neighbors by its average) and that 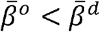, and also that *α*^o^ < *α^d^* **(Fig. S9).** Under these conditions, for the same *N_max_,* the following holds, 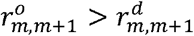, that is, the transition rate to gain new 3D neighbors is larger in the ordered case than in the disorder case. This is in fact the situation that we observed in the Voronoi tubular model when we estimated the value of the energy barrier to gain new neighbors: *α* and *β*(*s*) decrease as the CVT scale increases even though the surface tension energy is independent of the CVT scale (see **Fig. S9, Fig. 1,** and **Table S1).** However, it is possible to find large regions in terms of the values of 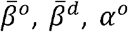, and *α^d^* where 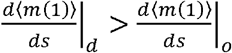. (**Fig. S11**). That is, the growth of 3D neighbors starting from the apical surface (i.e. *s* = 1) in the disorder case can be actually larger than that of the order case despite the fact that the transition rate to gain new 3D neighbors is smaller in the former.

Also, from Eq. (19), it is possible to infer, approximately, the expected behavior of 〈*m*(*s*)〉 = 〈*n*_3*D*_(*s*)〉 as follows. First, by performing a mean-field-like approximation, i.e., 〈*F*(*m*)〉 ≈ *F*(〈*m*〉),

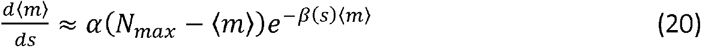

Second, assuming that *β*(*s*) < 1 (otherwise it is difficult to justify the observed presence of apico-basal intercalations),

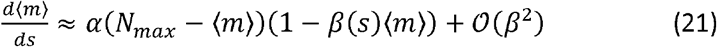

Eq. (21) is formally a logistic-like growth equation that can solved subjected to the condition 〈*m*(1)〉 = 6 (the average number of neighbors in the apical surface is 6).

We notice that in this case, the disorder levels of the wt and the mutant glands are similar. Consequently, the accumulation of 3D neighbors only depends on the transition rates, *r*_*m,m*+1_, that in turn are larger in the mutant background since the energy barrier decreases.

For finding the parameters of the Kolmogorov model, Eq. (19), that better fit *in silico* tubes and salivary glands we implemented an algorithm that solves, numerically, the set of equations defined by Eq. (19) and the normalization condition 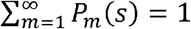 to obtain 〈*m*(*s*)〉 = ∑_*m*_*mP_m_*(*s*). Such algorithm minimizes the error between the curves 〈*m*(*s*)〉 obtained in the model and in experiments.

The values of the parameters obtained were further used to compare the predicted probability distribution of having *m* accumulated 3D neighbors for a given value of *s*: {*P_m_*(*s*)}. We evaluated the relative error of this prediction with respect to the actual distribution from data, 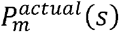, by computing 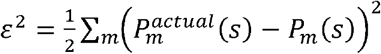. This quantity is normalized such that in case of the following situation of full disagreement between the distributions, 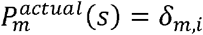 and *P_m_*(*s*) = *δ_m,j_* with *i* ≠ *j*, provides *ε*^2^ = 1 (i.e., 100% error).

### Quantitative characterization of spreading in neighbor exchange distributions between apical and basal surfaces

In order to characterize the spreading away from the diagonal in the neighbor exchange distributions between apical and basal surfaces, e.g., **Fig. 2A,** we followed the same approach used to quantify intrinsic noise during gene expression processes, see (Elowitz, 2002). Thus, 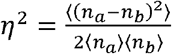 where 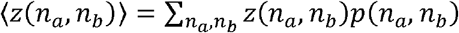; *z* representing any function of *n_a_* and *n_b_* and *p*(*n_a_, n_b_*) being the probability of neighbor exchange events. We point out that bins in the diagonal do not correspond necessarily to prismatic cells since a fraction of cells can conserved the polygonal class in apical and basal surfaces and yet undergo apico-basal intercalations.

### Statistical comparisons

The characteristics extracted from wildtype and mutant glands were compared by using a univariate statistical protocol **(Table S1).** This procedure allows to study if the data from two different groups of data follow a similar distribution: 1) we evaluated whether features values of these two kinds of glands presented normal distribution and similar variance using the Shapiro-wilk test and two-sample F-test respectively. 2) If data followed a normal distribution and had similar variance, we employed the two-tailed Student’s t-test. 3) In case, data presented a normal distribution but not equal variance we employed the two-tailed Welch test to compare means from both groups. 4) When data did not present normal distribution, we used the Wilcoxon test to compare medians from both groups.

In a different statistical analysis, we tested polygon distribution similarity from apical and basal surfaces of wildtype and mutant glands and V8 at *s_a_* = 1, *s_b_* = 1.75 and *s_b_* = 10 **(Table S1).** Following the guidelines from (Sánchez-Gutiérrez et al., 2016), we implemented chi-squared tests across all samples, being corrected for multiple testing using the method of Benjamini and Hochberg. To develop a more robust analysis, we used the distribution of 5-, 6- and 7-sided polygons due to the low presence (or inexistence) of the other kind of polygons (3-, 4-, 8-, 9-sided cells).

### Data and code availability

All the necessary materials to reproduce this study are available at Mendeley Data repository: DOI: 10.17632/gpz68wzhc2.1 and https://osf.io/nd5t6/.

