## Supplementary Information for "A quantitative biophysical principle to explain the 3D cellular connectivity in curved epithelia"

**SUPPLEMENTAL INFORMATION TEXT**

**Figure S1. A:** Percentage of scutoidal cells in the Voronoi model as a function of the surface ratio coordinate and the CVT scale. **B:** *In silico* and *in vivo* tubular epithelia satisfy a linear relation between the average number of 3D neighbors and the average number of apico-basal intercalations: $\left\langle n_{3D} \right\rangle=\left\langle i \right\rangle/2+6$. **C:** Average number of 3D neighbors per cell in the Voronoi model as a function of the surface ratio coordinate and the CVT scale.

**Figure S2**. Polygon distribution of salivary glands and Voronoi 8 (V8) *in silico* tubes: the error bar accounts for the standard deviation (𝑛 = 20 tubes). Green bars correspond to salivary gland and blue bars to V8 tubes. Lighter colors represent apical surfaces whereas darker are related to basal surfaces. The figure also shows the V8 case ($s_{b}=10$) that leads to an increase of topological disorder (larger variance of cell sidedness) in the basal surface.

**Figure S3.** (left) 3D histogram of neighbors in the apical and basal surface in the V8 model ($s_{b}=1.75$). (right, top) Cross-correlation in the V8 model ($s_{b}=1.75$) between average energy profiles along the apico-basal axis, $\left\langle E_{A} \right\rangle$ (dark grey), $\left\langle E_{V} \right\rangle$ (red), and $\left\langle E_{A} \right\rangle+\left\langle E_{L} \right\rangle$ (light grey), and the number of apico-basal intercalations, $\left\langle i \right\rangle$. (right, bottom) Cross-correlation between energy changes ($\left\langle E_{A} \right\rangle$ and $\left\langle E_{V} \right\rangle$) and the growth of intercalations along the apico-basal axis. Color code as in **Fig. 3B, D, E**.

**Figure S4.** **A:** “Poor get richer” principle in salivary glands and V8 tubes $s_{b}=1.75$ ($n=20$ tubes). The size of the circle accounts for the relative data count within each apical polygon class (numbers indicate the number of cells that gained 3D neighbors). The boxes indicate the $25\%-75\%$ percentile interval, black lines the mean values, gray lines the standard deviation, and the red dotted lines the statistical median. Cells with a smaller polygonal class are more prone to gain neighbors. (inset) Average net gain of neighbors with respect to the apical surface (i.e., black lines in main panel). **B:** "Poor get richer" in salivary glands and the V8 model ($s_{b}=1.75$): average net gain of neighbors, from basal to apical, as a function of the polygon class in the basal surface.

**Figure S5. A:** As a function of the longitudinal and transversal coordinates, this panel shows, schematically, an apico-basal intercalation from the point of view of a Voronoi diagram (black lines) and its topological dual, the Delaunay triangulation (gray lines). Small circles indicate the seeds of Voronoi cells and large circles show the Delaunay property graphically: nearest neighbors define triangles (cells’ seeds being their vertices) and circumscribed circles. In the apical surface, the nearest neighbors of the red seed are those seeds at the circumscribed red circle. Likewise, the nearest neighbors of the green seed are those seeds in the green circle. The distances between seeds along the longitudinal direction are preserved but transversal distances increase as the surface ratio increases from apical to basal. This anisotropy ultimately underlies apico-basal intercalations. Once an intercalation takes place (basal surface), the seeds in both circles are nearest neighbors of the red and the green seeds. **B:** Before the apico-basal intercalation shown in **A** occurs, the green seed is necessarily outside the circumscribed circle (otherwise it would be a nearest neighbor of the red seed). *Lemma 1* (**STAR Methods**) states that if the green seed is going to become a nearest neighbor of the red seed due to an apico-basal intercalation then it must be necessarily contained inside the green parabola. **C:** A consequence of *Lemma 1* is the so-called "poor get richer" principle. In this example, for a given fixed value of the surface ratio expansion, it is shown that the new neighbors of cell $P$ must be contained inside the parabolas $A$ and $B$: the more neighbors the cell has (from left to right), the smaller the region available to gain new neighbors (green shaded area). The probability, $\mathcal{P}$, of gaining new neighbors as a function of the actual number of neighbors, $n$, is then proportional to $S\left( A \right)+S\left( B \right)\leq S\left( \gamma\right)+S\left( \theta\right)$, where $S$ stands for the area and $\gamma$ and $\theta$ indicate the triangular sectors given by those angles. Since $S\left( \gamma\right)\sim S\left( \theta\right)\propto\frac{1}{n}$ then $\left( n \right)\propto\frac{1}{n}$ . An additional consequence of *Lemma 1* is that the 3D cellular connectivity gain decreases as the surface ratio increases: panels **D**-**F**. In this example, **D**-**F**, the perpendicular axis represents the longitudinal axis of tubes, while the horizontal axis accounts for the Cartesian projection of the transversal axis of radial sections. From left to right different radial sections are represented as $s$ increases (as indicated by the color gradient arrow: from light to dark blue). In **D** three Voronoi seeds that correspond to neighboring cells at the apical surface, $s=1$, define the triangle $ABC$. Panels **E** and **F** track changes in the neighboring relations (accumulated neighbors) of cell $A$ for two increasing values of $s$: $2$ and $4.5$ (panels **E** and **F** respectively). As shown in **D**, should a new neighboring cell, $D$, of cell $A$ appear due to an apico-basal intercalation, then its position must lie inside the vertical parabola defined by the points $A$, $B$ and $C$, but outside the circle that these points define (white region), (*Lemma 1*). Regions accessible to new neighbors are then coded by the green shading in **D**-**F**. As $s$ increases, see **E**, and the cells $A$, $B$, $C$, and $D$ become neighbors, then the parabolas and circles defined by $ABD$ and $ACD$ restrict the locations of future nearest neighbors. This idea is further reinforced in panel **F**: winning neighbor $E$ set additional limits to the accessible locations of new neighbors. Thus, the potentiality of a connectivity gain by cell $A$ due to apico-basal intercalations diminishes as the surface ratio increases and eventually becomes null: the number of 3D neighbors of a cell is bounded (**STAR Methods**).

**Figure S6.** Applicability of the biophysical model for different cases of the Voronoi model in terms of the CVT scale. Color codes and legends as in **Fig. 4D, E**.

**Figure S7. A:** Levels of E-cadh protein expression along the cell apico-basal axis in wildtype (green) and ∆Ecad (orange) glands ($n=30$ cells per phenotype) (**STAR Methods**). The solid lines stand for the mean and the bands for the standard deviation. **B-F:** Statistical comparison between wildtype ($n=20$ glands, green) and ∆Ecad ($n=10$ glands, orange) glands properties: cell volume, $s_{b}^{*}$ (effective basal surface ratio), proportion of scutoids, $n_{3D}$ (number of 3D neighbours) and $i$(number of apico-basal intercalations), respectively (see **Table S1**). In each panel, the solid circles represent the average value of the corresponding feature for each gland, long horizontal black lines denote the mean value, and small ones the standard deviation. The protocol for the statistical quantification of data is described in **STAR Methods**. Significant difference $p<0.05$ *, $p<0.01$ **, and $p<0.001$ ***; No significant difference, ns.

**Figure S8. A:** Density plot of the 3D distribution of neighbor exchanges between apical and basal surfaces as a function of the number of neighbors in apical, $n_{a}$, and basal, $n_{b}$, surfaces (as in **Fig. 2A**) in mutant salivary glands. **B:** “Poor get richer” principle in ∆Ecad salivary glands: average net gain of neighbors, from apical to basal, as a function of the polygon class in the apical surface. **C:** Average profiles of the number of apico-basal intercalations, $\left\langle i \right\rangle$, average lateral area, $\left\langle A \right\rangle$, and cellular volume fluctuations, $\sigma_{V}^{2}$ in *in vivo* mutant tubes as a function of the apico-basal coordinate, $s$. **D-E:** Cross-correlation analysis between energy and intercalation profiles: same as **Fig. 3D, E**. ($n=10$ glands).

**Figure S9.** Energy cost to gain additional 3D neighbors per cell (top) and the corresponding “bare” transition rate (bottom) in the Voronoi tubular models (V1 to V10, $s_{b}=10$) as estimated by the Kolmogorov model.

**Figure S10.** A scutoid solid can be simplified as a collection of vertexes (circles) that can be connected by straight edges. The green circle accounts of an apico- basal intercalation point. Alternatively, we can represent the solid as a connected plane graph. In both cases the Euler characteristic is 2.

**Figure S11.** In the parameter space $\left\{ \bar{\beta}^{o},\bar{\beta}^{d},\alpha^{d}/\alpha^{o} \right\}$ (see **STAR Methods**) it is possible to find large regions where the disorder facilitates the accumulation of 3D neighbors (green shaded region) even if the transition rates to gain 3D neighbors in more ordered configurations are larger. The red shaded region indicates in the same parameter space the region where both the transition rates and the accumulation of 3D neighbors are larger in an ordered configuration.

**Table S1.** Tab 1 (**Glands properties**), shows some glands properties and the statistical differences after comparing wildtype and mutant gland features, Tab 2 (**Stats polygon distributions**) shows the statistical comparison between apical and basal surfaces of glands and V8 surfaces: $s_{a}=1$, $s_{b}=1.75$ and $s_{b}=10$, Tab 3 (**Kolmogorov**) shows the independent fitting parameter values ($\alpha$, $\beta$, and $\left\langle N_{max} \right\rangle$), Tab 4 (**Spreading**) shows the spreading values of the 3D histograms of neighbors exchange between basal and apical surfaces (**STAR Methods**).
